# Apoptosis inhibition reprograms alveolar myofibroblasts toward ductal myofibroblasts

**DOI:** 10.1101/2025.05.26.654588

**Authors:** Maria Jose Gacha-Garay, Hui Liu, Scott E Evans, Tingting W Mills, Jichao Chen

## Abstract

The epithelial tree of the lung is shaped proximo-distally by airway smooth muscle cells (ASMCs), ductal myofibroblasts (DMFs), and, transiently, alveolar myofibroblasts (AMFs). Lineage tracing and snapshot imaging suggest the clearance of AMFs via apoptosis post-alveologenesis, although definitive evidence is lacking. Here, we generate an inducible BCL2 overexpression mouse allele to inhibit AMF apoptosis. Using three independent Cre drivers and single-cell RNA-seq, we show that BCL2-rescued AMFs persist around distal alveolar ducts and alveoli and, unexpectedly, mature toward DMFs. Both normal DMFs and rescued DMF-like cells upregulate contractile proteins in a house dust mite-induced asthma model. Our findings demonstrate the apoptotic clearance of AMFs, as well as fate plasticity and pathophysiological convergence of lung mesenchymal cells of the epithelial axis.

## INTRODUCTION

Cells change fates and states during development to form diverse cell types and upon perturbation to ensure tissue repair or to drive degenerative, fibrotic, or tumorigenic pathology. This cellular plasticity—the ability of cells to interconvert—often can be deduced from spatial proximity, transcriptional similarity, and developmental lineage. The immune lineage hierarchy represents a textbook example owing to transplantation assays and myriad cell surface markers^1^. Research on plasticity within epithelial, endothelial, and neuronal lineages has benefited from their stereotypic arrangement around and along lumens^2–7^. However, the mesenchymal lineage, especially the variety of fibroblasts with non-discriminating space-filling cell morphology, is not well defined to ascertain cellular plasticity, despite its importance in diseases such as cancer and fibrosis^8–13^.

Using scRNA-seq and 3D imaging, we have classified mouse lung mesenchymal cells into three axes: vascular, epithelial, and interstitial, each organized around the vascular and epithelial trees and the space in between^14^. From proximal to distal, the vascular axis includes vascular smooth muscle cells (VSMCs) and pericytes; the epithelial axis includes airway smooth muscle cells (ASMCs), ductal myofibroblasts (DMFs), and alveolar myofibroblasts (AMFs, which include secondary crest myofibroblasts (SCMF)); while the interstitial axis includes adventitial fibroblasts (AF2s, formerly proximal interstitial fibroblasts) and alveolar fibroblasts (AF1s, formerly distal interstitial fibroblasts). The spatial and transcriptional relatedness among cells within each axis suggests potential cellular plasticity—as exemplified by the ability of pericytes to become vascular smooth muscle cells^15,16^—a phenomenon we will further investigate within the epithelial axis in this study.

Of the epithelial axis, ASMCs, DMFs, and AMFs are all contractile and confine the airways, alveolar ducts, primary and secondary septa^14,17–19^. Extending from conducting airways, the alveolar ducts are multi-branch generational tubes lined with alveolar epithelial cells. Their distal ends open into alveolar sacs and thus correspond to alveolar entrance rings and primary septa^20,21^. The associated DMFs are marked by HHIP and CDH4, but not PDGFRA, whose high expression marks AMFs^14,22^. AMFs, particularly their contractility, are essential for secondary septation to increase alveolar surface area and constitute nearly half of lung mesenchymal cells at the peak of neonatal alveologenesis before disappearing^14,23,24^. *Pdgfra^CreER^*and Myh11-CreER lineage-tracing experiments rule out their reprogramming into other lung mesenchymal cell types, while apoptosis markers, including cleaved Caspase-3 and pyknotic nuclear profiles, peak in AMFs at P12-P13 just before their disappearance ^14,25,26^. However, definitive evidence demonstrating apoptosis as the primary mechanism for AMF clearance is still lacking.

Developmental apoptosis eliminates excess cells to shape tissues, as seen in digit separation, and refines function, as observed in neuronal pruning and immune tolerance^27–31^. Studying apoptosis in vivo is inherently challenging due to its asynchronous nature and rapid phagocytic clearance. In this study, leveraging well-defined pro-apoptotic and pro-survival molecular pathways, we generate a Cre- and Flp-inducible BCL2 overexpression allele to tilt the balance toward cell survival. Using three independent genetic drivers targeting AMFs, we rescue AMFs via BCL2 overexpression and reveal their plasticity toward the DMF fate, as well as pathophysiological convergence toward a DMF response to house dust mite-induced allergic inflammation.

## RESULTS

### An inducible BCL2 overexpression model inhibits apoptosis in a cell-specific manner

To cumulatively inhibit transient, asynchronous apoptosis in a cell population, we opted for cell-type-specific, continuous expression of a pro-survival gene BCL2, known to counteract the pro-apoptotic machinery in immune cells ^32–35^. Accordingly, we engineered a Flp and Cre-dependent FLAG-tagged BCL2^36^ allele in the *Rosa* locus (Figure 1A). The FRT-flanked transcriptional stop cassette enabled intersectional spatiotemporal control but was unnecessary for this study and was therefore removed by crossing to *Rosa^Flp^* ^37^. For simplicity, we named the original double-conditional allele as *Rosa^BCL2+F^*and the Flp-recombined allele as *Rosa^BCL2^* (Figure 1A and 1B).

**Figure 1.**
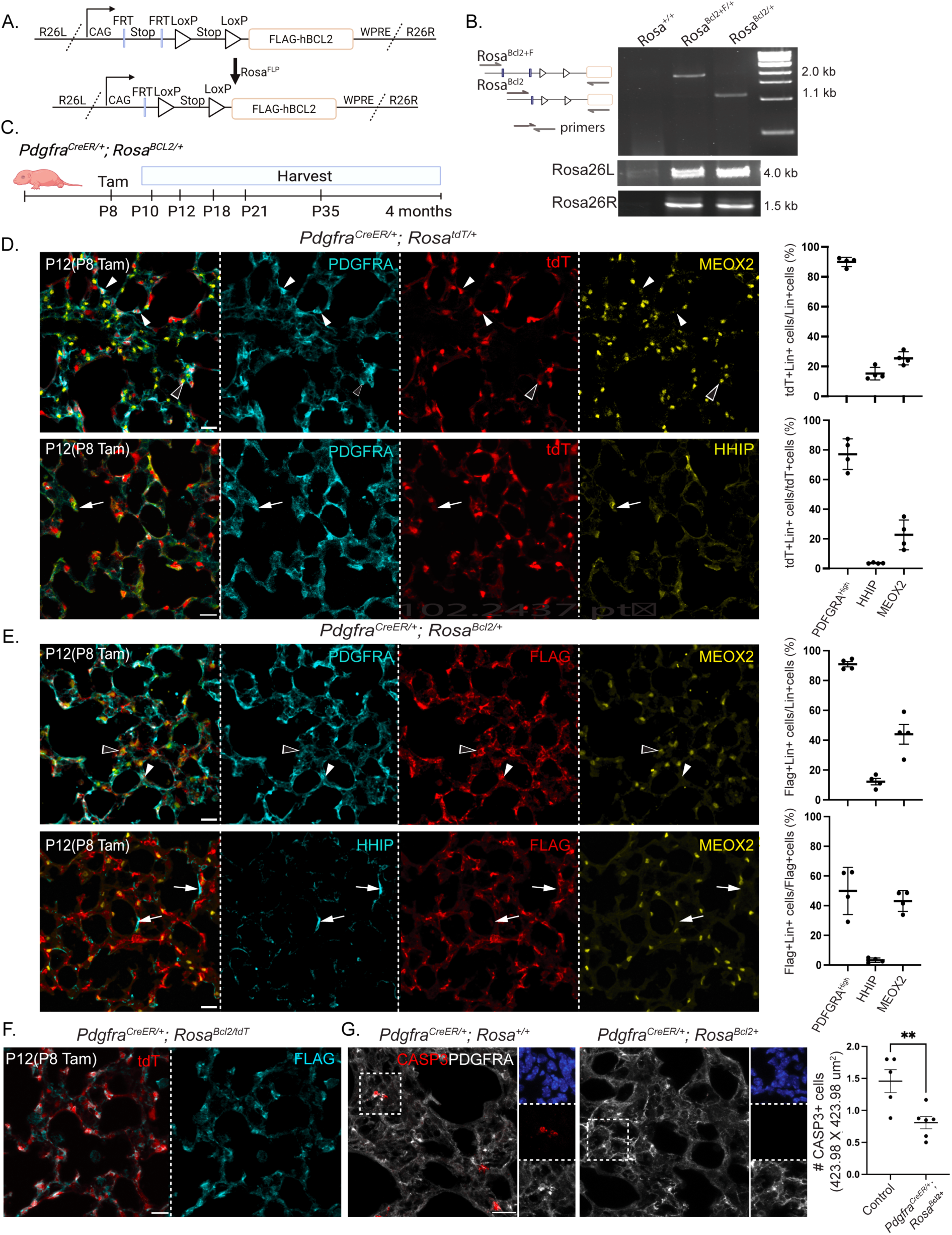
An inducible BCL2 overexpression model inhibits apoptosis in a cell-specific manner. A. Diagram of original double-conditional allele (*Rosa^BCL2+F^)* showing FLAG-tagged human BCL2 driven by a constitutive promoter (CAG) and preceded by two stop transcriptional cassettes flanked by FRT and LoxP sites, respectively, followed by a stabilizing Woodchuck hepatitis virus posttranscriptional regulatory element (WPRE) and inserted between left and right homology arms targeting the *ROSA26* locus (R26L and R26R). Removal of the FRT-flanked stop cassette with *Rosa^Flp^* generates the Cre-dependent *Rosa^BCL2^*allele. B. PCR genotyping of *Rosa^BCL2+F^* and *Rosa^BCL2^* alleles validates removal of FRT-flanked stop cassette and insertion in the *ROSA26* locus. C. Experimental design showing induction of BCL2 overexpression driven by *Pdgfra^CreER^* with tamoxifen (Tam) injection at P8 and evaluation during alveologenesis (P10-P21) and adulthood (P35 and 4 months). D. Immmunostaining and quantification of *Pdgfra^CreER/+^; Rosa^tdT/+^*lungs showing efficient labeling of PDGFRA^high^ AMFs (arrowhead), moderately MEOX2+ AF1s (open arrowhead), and occasionally HHIP+ DMFs (arrow: unlabeled DMFs). Lin: lineage. tdT level in AF1s is lower than that in AMFs and DMFs. AF2s in bronchovascular bundles are not included. Each symbol represents a mouse; average of three images of 424 x 424 um^2^ per mouse; error bar represents s.e.m. (see Table S1 for raw data; same for all figures unless noted otherwise). E. Immunostaining and quantification of *Pdgfra^CreER/+^; Rosa^BCL2/+^*lungs showing comparable labeling efficiency and specificity with the *Rosa^tdT^* reporter. See D for details. F. Immunostaining of *Pdgfra^CreER/+^; Rosa^BCL2/tdT^* lungs, representative of at least 3 mice, showing FLAG and tdT colocalization. G. Immunostaining and quantification of *Pdgfra^CreER/+^; Rosa^BCL2/+^* and control lungs showing that BCL2 overexpression reduces developmental AMF apoptosis (CASP3+PDGFRA^high^). Asterisk: p=0.0087 (Mann-Whitney test). All scale bars: 20 um.

To validate our BCL2 allele in vivo, we activated *Rosa^BCL2^*throughout the lung epithelium using *Shh^Cre^* ^38^ and confirmed FLAG-BCL2 expression in nearly all NKX2-1+ epithelial cells, correctly colocalizing with the mitochondrial marker TOMM22^39^ (Figure S1A). *Shh^Cre^* also targeted the posterior digits in the limb, leading to webbing specifically between the fourth and fifth digits in our BCL2 overexpression mutant (Figure S1B). This phenotype recapitulated the classic example of failed developmental apoptosis, similar to that observed in the *Bax*/*Bak* double mutant^31^. Additionally, we used Sox2-Cre to activate *Rosa^BCL2^* in the epiblast, thereby affecting all somatic tissues^40^. The resulting mutant had cleft palate, sacral spina bifida, and curled tail, resembling the phenotype of triple mutants in the pro-apoptotic genes *Bax*, *Bak*, and *Bok*^27^ (Figure S1C). Thus, our BCL2 model inducibly blocks developmental apoptosis.

### *Pdgfra^CreER^*-driven BCL2 overexpression inhibits apoptosis in AMFs

As AMFs and DMFs were distinguishable by PDGFRA-high and HHIP/CDH4 expression, respectively, by postnatal day (P) 7, and noticeable AMF apoptosis started from P10^14,26,41^, we induced BCL2 overexpression with *Pdgfra^CreER^* at P8 (Figure 1C). Since MEOX2+ AF1s also expressed PDGFRA, albeit at a lower level^14,42^, we assessed the efficiency and specificity of *Pdgfra^CreER^* using a *Rosa^tdT^* reporter (Figure 1D). Four days post-tamoxifen injection, 91% of PDGFRA-high AMFs, 15% HHIP+ DMFs, and 25% MEOX2+ AF1s were tdT+, while 73%, 3%, and 23% of tdT+ cells were AMFs, DMFs, and AF1s, respectively (Figure 1D). Similarly, for the BCL2 allele in the same *Rosa* locus, 91% of AMFs, 10% of DMFs, and 45% of AF1s were FLAG+, while 49%, 3%, and 44% of FLAG+ cells were AMFs, DMFs, and AF1s, respectively (Figure 1E). AF2s in the bronchovascular bundles were not counted. The specificity percentages within tdT+ cells did not sum to 100% due to the necessity of separate immunostaining panels for the three cell types. Native tdT fluorescence was dimmer in AF1s compared to AMFs and DMFs, potentially due to delayed recombination from lower driver expression and/or lower *Rosa* locus activity in AF1s. However, this discrepancy was not observed for FLAG, which was amplified through immunostaining (Figure 1E). Finally, tdT and FLAG-BCL2 co-expressed in nearly all recombined cells in *Rosa^tdT/BCL2^*lungs (Figure 1F).

Consistent with the observed digit webbing and body malformation phenotypes (Figure S1B and S1C), *Pdgfra^CreER/+^; Rosa^BCL2/+^* lungs had fewer PDGFRA-high cells marked by cleaved Caspase-3 (Figure 1G). This reduction was likely underestimated, as apoptotic cells were expected to rapidly lose their cell-type markers, including PDGFRA. Additionally, any inhibition of apoptosis in dying cells at a given time point would accumulate due to the broad, continuous expression of BCL2. Taken together, *Pdgfra^CreER^* activation at P8 efficiently targets AMFs, moderately targets AF1s, and minimally targets DMFs, which is relevant for phenotype interpretation below. BCL2 overexpression effectively inhibits AMF apoptosis.

### Persistent AMFs are reprogrammed toward the DMF fate around distal alveolar ducts and alveoli

Consistent with our previous findings^14^, in P35 lungs labeled with tamoxifen injection at P8, nearly all tdT+ cells (92%) in the control lungs were MEOX2+ AF1s, as expected from dual labeling of AMFs and AF1s by *Pdgfra^CreER^* and subsequent apoptotic clearance of labeled AMFs (Figure 2A). However, in BCL2 lungs, only 56% of tdT+ cells were AF1s, with the remainder named “persistent AMFs” because they escaped apoptosis owing to BCL2 overexpression (Figure 2A). Importantly, the number of MEOX2+ cells in control and BCL2 lungs remained unchanged, indicating a lack of BCL2-sensitive apoptosis in AF1s (Figure 2A).

**Figure 2.**
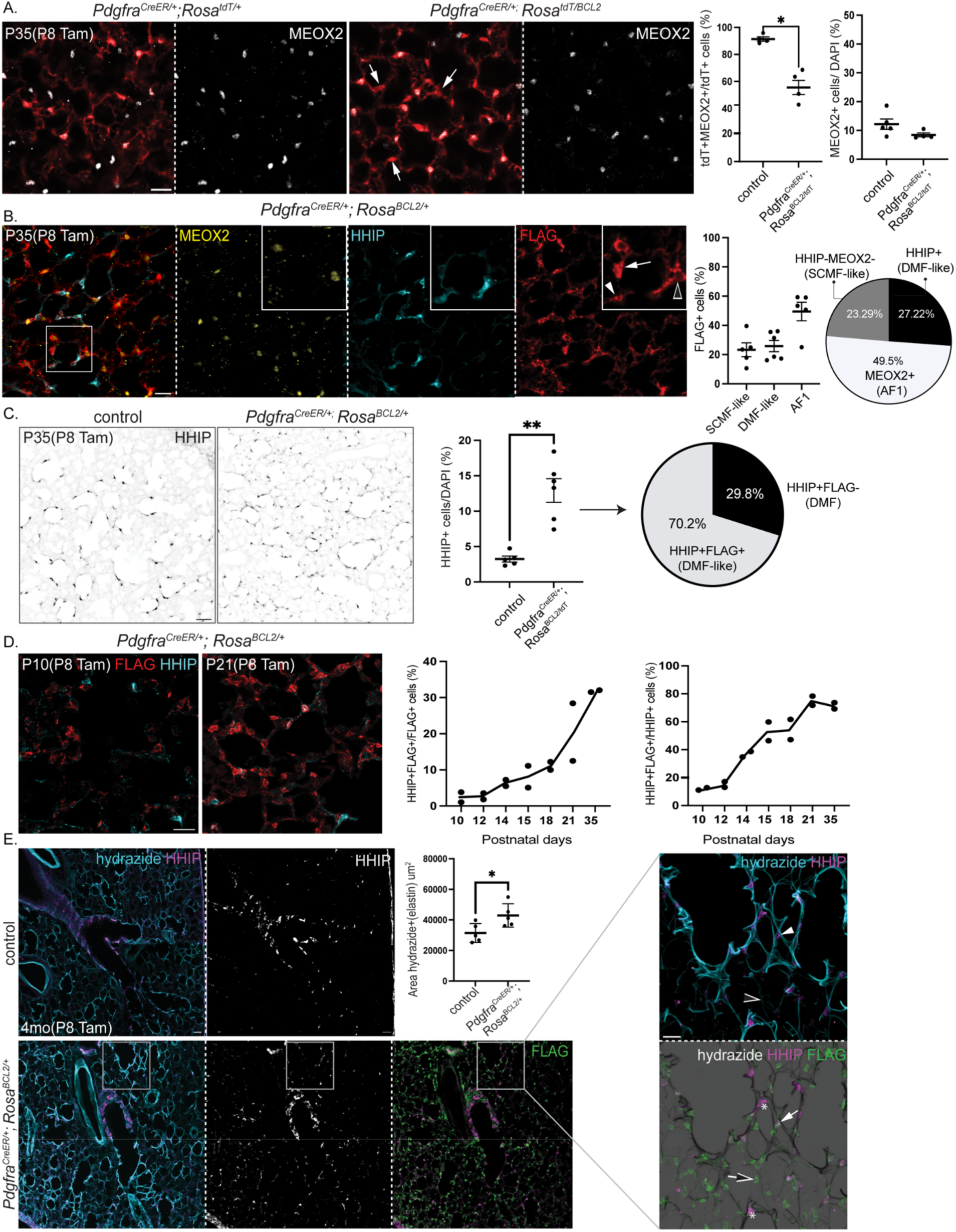
Persistent AMFs are reprogrammed toward the DMF fate around distal alveolar ducts and alveoli. A. Immunostaining and quantification of mature *Pdgfra^CreER/+^; Rosa^tdT/+^* control and *Pdgfra^CreER^; Rosa^BCL2/tdT^* lungs showing more persistent MEOX2-tdT+ cells (arrow) in the BCL2 lung. The number of AF1s (MEOX2+) is unchanged (p=0.0635; Mann-Whitney test). Scale bar = 20 um (same for all panels unless noted otherwise). B. Immunostaining and quantification of mature *Pdgfra^CreER/+^; Rosa^BCL2/+^* lungs showing three FLAG+ populations: expected AF1s (MEOX2+; open arrowhead) and aberrant DMF-like (HHIP+; arrowhead) and SCMF-like (MEOX2-HHIP-; arrow) cells. C. Immunostaining and quantification of mature *Pdgfra^CreER/+^; Rosa^BCL2/+^* and control lungs showing more HHIP+ cells in BCL2 lungs (p=0.0043; Mann-Whitney test). Pie chart: in BLC2 lungs, 70.2% HHIP+ cells are DMF-like cells. D. Immunostaining and quantification of *Pdgfra^CreER/+^; Rosa^BCL2/+^*lungs over time showing an increasing proportion of FLAG cells becoming DMF-like (HHIP+) and an increasing proportion of HHIP+ cells becoming lineage traced post developmental apoptosis of AMFs. E. Immunostaining and quantification of increased elastin fiber area (p=0.0317; Mann-Whitney test) in adult *Pdgfra^CreER/+^; Rosa^BCL2/+^* and control lungs showing expansion of HHIP+ cells from proximal to distal alveolar ducts. DMF (FLAG-; asterisk) and DMF-like cells (FLAG+; arrow) are outlined by thick hydrazide-stained elastin fibers (arrowhead) while SCMF-like (FLAG+HHIP-; open arrow) are outlined by thin fibers (open arrowhead) upon BCL2 expression. Scale bar: 50 um

We then sought to define the persistent AMFs with our published mesenchymal cell markers^14^. Among FLAG-BCL2 expressing cells, 50% were MEOX2- and thus persistent AMFs; the slight percentage discrepancy with the tdT reporter likely arose from experimental and allele variations. These cells did not express markers of the vascular axis (PDGFRB and NOTCH3) nor high levels of PDGFRA, suggesting that they were no longer bona fide AMFs (Figure S2A, S2B). Unexpectedly, half of persistent AMFs (27% of FLAG+ cells) ectopically expressed DMF markers HHIP and CDH4, which we named “DMF-like cells” (FLAG+HHIP+), whereas the rest (23% of FLAG+ cells) were called “SCMF-like cells” (FLAG+HHIP-MEOX2-)—a classification further supported by scRNA-seq below (Figure 2B, S2C). As a result, the proportion of HHIP+ cells rose from 3% in control lungs to 16% in BCL2 lungs, such that 70% of the HHIP+ cells in BCL2 lungs were FLAG+ DMF-like cells (Figure 2C). Persistent AMFs (i.e., DMF-like and SCMF-like cells)—notably, even normal DMFs—no longer expressed the contractile genes SMA and TAGLN, suggesting that the contractility in the alveolar region required for neonatal alveologenesis diminished in the mature lung (Figure S2B).

Next, we examined the spatiotemporal distribution of persistent AMFs. Temporally, DMF-like cells (FLAG+HHIP+), quantified as a proportion of all HHIP+ or FLAG+ cells, increased over time, coinciding with AMF clearance (Figure 2D). Spatially, in P35 and 4-month-old control lungs, HHIP+ DMFs were confined to proximal alveolar ducts, extending 2-3 branch generations from the terminal bronchioles (Figure 2E). This raised the possibility that cells surrounding more distal alveolar ducts were AMFs, instead of DMFs, and were lost following neonatal alveologenesis. In contrast, in P35 and 4-month-old BCL2 lungs, DMF-like cells surrounded distal alveolar ducts with thick elastin fibers, while SCMF-like cells were positioned further distally around the alveoli with thin or no elastin fibers, although infrequently they could be juxtaposed (Figure 2E and S3). Elastin fibers, stained with hydrazide^43^, increased in BCL2 lungs; however, mean linear intercept measurements of alveolar morphology and FlexiVent measurements of lung mechanics were unaffected except for a small decrease in static compliance (Figure 2E and S4).

Our *Pdgfra^CreER^* driver experiments demonstrate that AMFs surrounding distal alveolar ducts and alveoli are normally cleared following neonatal alveologenesis. These cells are selectively targeted by *Pdgfra^CreER^*and persist upon BCL2 overexpression as DMF-like and SCMF-like cells until at least 4 months of age.

### Single-cell RNA-seq identifies persistent AMFs as DMF-like and SCMF-like cells

To confirm and fully define persistent AMFs, we conducted scRNA-seq on lungs from P38 *Pdgfra^CreER/+^; Rosa^BCL2/+^* and control *Pdgfra^CreER/+^; Rosa^tdT/+^*mice induced with tamoxifen at P8. Lungs from male and female mice were distinguishable by *Xist* expression and were comparable (Figure S5A). We computationally separated the *Col1a1*+ mesenchymal lineage from *Cdh1*+ epithelial, *Icam2*+ endothelial, and *Ptprc*+ immune lineages, then re-clustered them to identify mesenchymal cell types along the vascular (VSMCs and pericytes), epithelial (ASMCs and DMFs), and interstitial (AF2s and AF1s) axes, as well as mesothelial cells (Figure 3A and S5A). While AMFs were absent in control lungs due to apoptotic clearance, the BCL2 lungs contained an ectopic cell population clustering near DMFs, with an abundance consistent with persistent AMFs (Figure 3A). These cells were identified as persistent AMFs also due to their positivity for WPRE—a 3’-UTR regulatory sequence present in both tdT and BCL2 alleles that functioned as a lineage marker. While DMFs in the control lung were rarely WPRE+, reflecting mistargeting of *Pdgfra^CreER^*, some DMFs in the BCL2 lungs were WPRE+, suggesting full conversion of a subset of persistent AMFs to DMFs. Persistent AMFs were negative for *Meox2*, a marker of AF1s and AF2s, which were also WPRE+ and thus part of the *Pdgfra^CreER^*lineage (Figure 3A). The possibility that BCL2 overexpression rescued AF1s and AF2s and converted them into the ectopic population was later ruled out through experiments using genetic drivers that did not target AF1s or AF2s.

**Figure 3.**
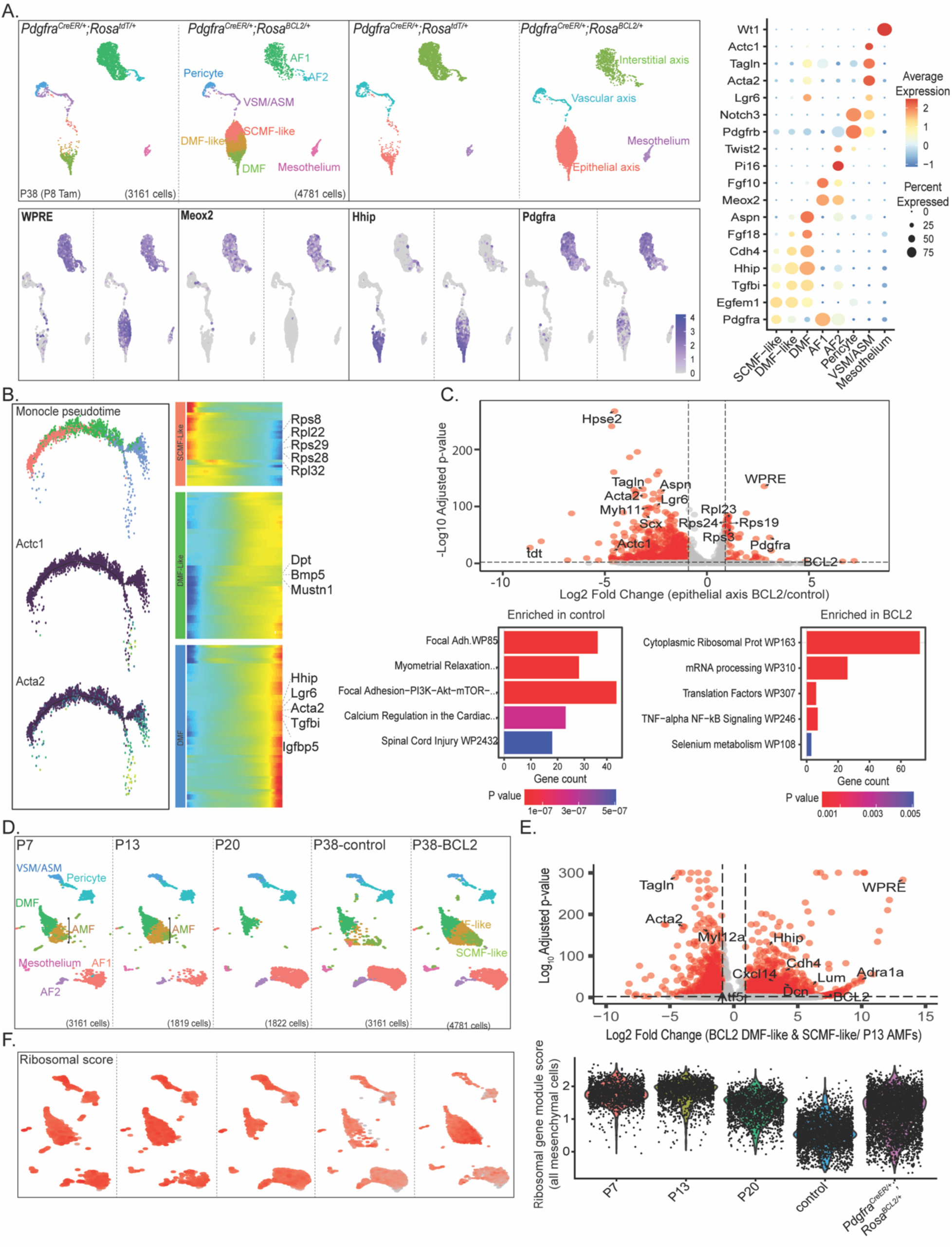
Single-cell RNA-seq identifies persistent AMFs as DMF-like and SCMF-like cells. A. Top: UMAPs of lung mesenchymal cells color-coded by cell type (left) and axis (right) identify two clusters—DMF-like and SCMF-like—that are present only in *Pdgfra^CreER/+^; Rosa^BCL2/+^* lungs (1 male and 1 female mice in each genotype). Bottom: Feature plots showing the lineage marker WPRE in AF1/2, DMF-like, and SCMF-like cells. Persistent AMFs unique to BCL2 lungs are negative for Meox2 and contain two subsets with opposing gradients of Pdgfra and Hhip, corresponding to SCMF-like and DMF-like cells in Fig. 2. Right: Dot plots of mesenchymal cell type markers showing that SCMF-like and DMF-like cells share AMF/DMF markers (Egfem1 and Tgfbi) but diverge for an AMF marker (Pdgfra) and DMF markers (Hhip, Cdh4, and Aspn). Fgf18 is higher in DMF-like cells than SCMF-like cells. B. Left: Monocle analysis of color-coded (as in A) mesenchymal cells of the epithelial axis from BCL2 lungs, showing a transcriptomic continuum from SCMF-like to DMF-like and then DMF cells. A bifurcation within DMFs, marked by increased contractile genes Actc1 and Acta2, corresponds to ASMCs or intermediate ASMC-DMF cells. Right: Trajectory heatmap showing ribosomal genes in SCMF-like cells, consistent with an immature state, and DMF markers in DMF-like cells. C. Top: Volcano plot comparison of mesenchymal cells of the epithelial axis from control and BCL2 lungs showing increased ribosomal genes in BCL2 lungs and decreased contractile genes and DMF markers, driven by SCMF-like cells. Bottom: Geno ontology analysis shows pathway enrichment in control (left) and BCL2 (right) lungs. See Table S2 for raw data (same for other scRNA-seq analyses). D. Integrated UMAPs of mesenchymal cells from normal lungs during alveologenesis and mature control and BCL2 lungs, showing that SCMF-like and DMF-like cells cluster with AMFs at P7 and P13. E. Volcano plot comparison of persistent AMFs (SCMF-like and DMF-like) from BCL2 lungs and AMFs from P13 lungs, showing enrichment of DMF and contractile markers, respectively. F. Feature (left) and violin (right) plots showing that the expression of the Seurat module score of ribosomal genes enriched in BCL2 lungs is higher in postnatal lungs compared to mature lungs (P20 and P38-control).

We further subdivided the ectopic population into two clusters, SCMF-like and DMF-like, based on opposing gradients and resulting cluster-specific enrichment of *Pdgfra* and *Hhip*, markers of AMFs and DMFs, respectively (Figure 3A). Both clusters expressed *Egfem1* and *Tgfbi*, shared markers of AMFs and DMFs (Figure 3A). Despite differences in measuring RNA versus protein and assay sensitivity, the percentages of AF1s (*Meox2*+), SCMF-like cells (*Hhip*−), and DMF-like and DMF cells (*Hhip*+) carrying the WPRE lineage marker in scRNA-seq (18%, 33%, and 43%, respectively), were comparable to the quantification of FLAG+ cells from imaging (Figure 2B). Compared to SCMF-like cells that had higher *Pdgfra*, DMF-like cells had expected enrichment of DMF markers, including *Aspn* and *Lgr6* (Figure 3A and S5B). Similarly, compared to DMF-like cells, DMFs had an even higher expression of *Aspn* and *Lgr6*, along with increased expression of contractile genes such as *Acta2* and *Tagln*, likely due to the presence of co-clustered ASMCs or intermediate ASMC-DMF cells (Figure 3A and S5B). A Monocle2^44^ trajectory analysis positioned DMFs, DMF-like cells, and SCMF-like cells along a differentiation spectrum, with distinct gene enrichment patterns, including ribosomal gene expression in SCMF-like cells (Figure 3B). A subset of Actc1+Acta2+ cells branched off DMFs, supporting co-clustering of ASMCs or intermediate ASMC-DMF cells (Figure 3B). A comparison of alveolar mesenchymal cells of the epithelial axis (DMFs and persistent AMFs) between control and BCL2 lungs confirmed that BCL2 lungs were enriched in ribosomal genes and mRNA processing and translation pathways. In contrast, even though control lungs had fewer total cells, they showed higher expression of DMF and ASMC markers and pathways of extracellular matrix and contractility, as only a subset of persistent AMFs were reprogrammed toward DMFs (Figure 3C).

Since persistent AMFs were rescued from apoptosis by BCL2 overexpression, we integrated scRNA-seq data from P38 control and BCL2 lungs with our previously published developmental time course data from P7, P13, and P20 normal lungs. On UMAP, persistent AMFs in the BCL2 lung aligned with developmental AMFs at P7 and P13 (Figure 3D). Comparison of P13 AMFs and P38 persistent AMFs revealed that, upon rescue by BCL2, AMFs upregulated DMF markers, including *Hhip* and *Cdh4*, supporting their reprogramming toward the DMF fate. Upregulated genes also included *Cxcl14*, *Atf5*, and *Adra1a*, which were typically absent and might indicate cellular stress resulting from forced survival^45–47^ (Figure 3E). Concurrently, they downregulated contractile genes, including *Talgn* and *Acta2*, indicating a deactivation of contractility necessary for neonatal alveologenesis—consistent with immunostaining (Figure S2B). Furthermore, ribosomal genes upregulated in persistent AMFs (Figure 3F) normally decreased as neonatal lungs matured, supporting a developmental origin for persistent AMFs (Figure 3D).

Although AMFs and DMFs had distinct proximo-distal locations and functions, they likely originated from progenitors recruited into the epithelial axis, as demonstrated for ASMCs in previous studies^22,48^. Consequently, AMFs captured at specific developmental time points, such as P7 or P13, exhibited heterogeneity, with a subset clustering near DMFs and potentially differentiating toward them—a process terminated by apoptosis. Supporting this possibility, published markers of immature (*Zmat5*, *Col24a1*, *Tenm4*, and *Lef1*), intermediate (*Wnt11* and low levels of *Tgfbi*, *Mustn1*, *Hhip*, *Thbs1*, and *Wnt5a*), and mature (*Lgr6*, *Gja1*, *Fhod3*, and high levels of *Hhip*, *Aspn*, *Lum*, and *Wnt5a*) myofibroblasts aligned with the two subpopulations of AMFs and the DMFs at P7 and P13, as well as SCMF-like cells, DMF-like cells, and DMFs in the P38 BCL2 lung (Figure S5C).

Finally, comparison of untargeted DMFs over time revealed that DMF maturation involved upregulation of extracellular matrix genes, including Lum and Dcn, and downregulation of contractile genes, including *Acta2* and *Myh11* (excluding the co-clustered ASMCs or ASMC-DMF intermediate cells), suggesting a switch from contractile to matrix-synthesis function (Figure S5D and S5E). No significant transcriptomic changes were detected in other cell lineages of the BCL2 lung, supporting cell-autonomous phenotypes of BCL2 overexpression (Figure S6A). CellChat analysis identified more ligand-receptor interactions involving DMFs than DMF-like and SCMF-like cells (Figure S6B and S6C). Notable differential signaling toward DMF-like versus SCMF-like cells included Semaphorin and Bmp signaling pathways, possibly promoting the DMF fate (Figure S6D and S6E).

Therefore, the contractility of DMFs and AMFs, essential for shaping the expanding alveolar ducts and alveoli during neonatal alveologenesis, is deactivated through transcriptional downregulation in DMFs and apoptotic clearance in AMFs. DMFs upregulate extracellular matrix genes, likely contributing to structural stability. BCL2-rescued AMFs mimic DMFs by reducing contractile gene expression, and a subset of them, DMF-like cells, upregulate matrix-synthesis genes.

### Two additional Cre drivers recapitulate the persistent AMF phenotype

To overcome the limitation of *Pdgfra^CreER^* targeting AF1s and AF2s in addition to AMFs, we used Myh11-CreER, which we previously demonstrated to specifically target contractile cells (VSMCs, ASMCs, DMFs, and AMFs) and pericytes while excluding AF1s and AF2s^14^ (Figure 4A and S7). In P21 Myh11-CreER; *Rosa^BCL2/+^* lungs, we again observed an expansion of HHIP+FLAG+ cells (25% of FLAG+ cells) to distal alveolar ducts, although we could not distinguish DMF-like cells from DMFs since both were lineage-labeled by Myh11-CreER (Figure 4B and 4C). Around the alveoli, in addition to the expected PDGFRB+FLAG+ pericytes (32% of FLAG+ cells), we identified PDGFRB-HHIP-FLAG+ cells (42% of FLAG+ cells), which likely corresponded to SCMF-like cells (Figure 4C).

**Figure 4.**
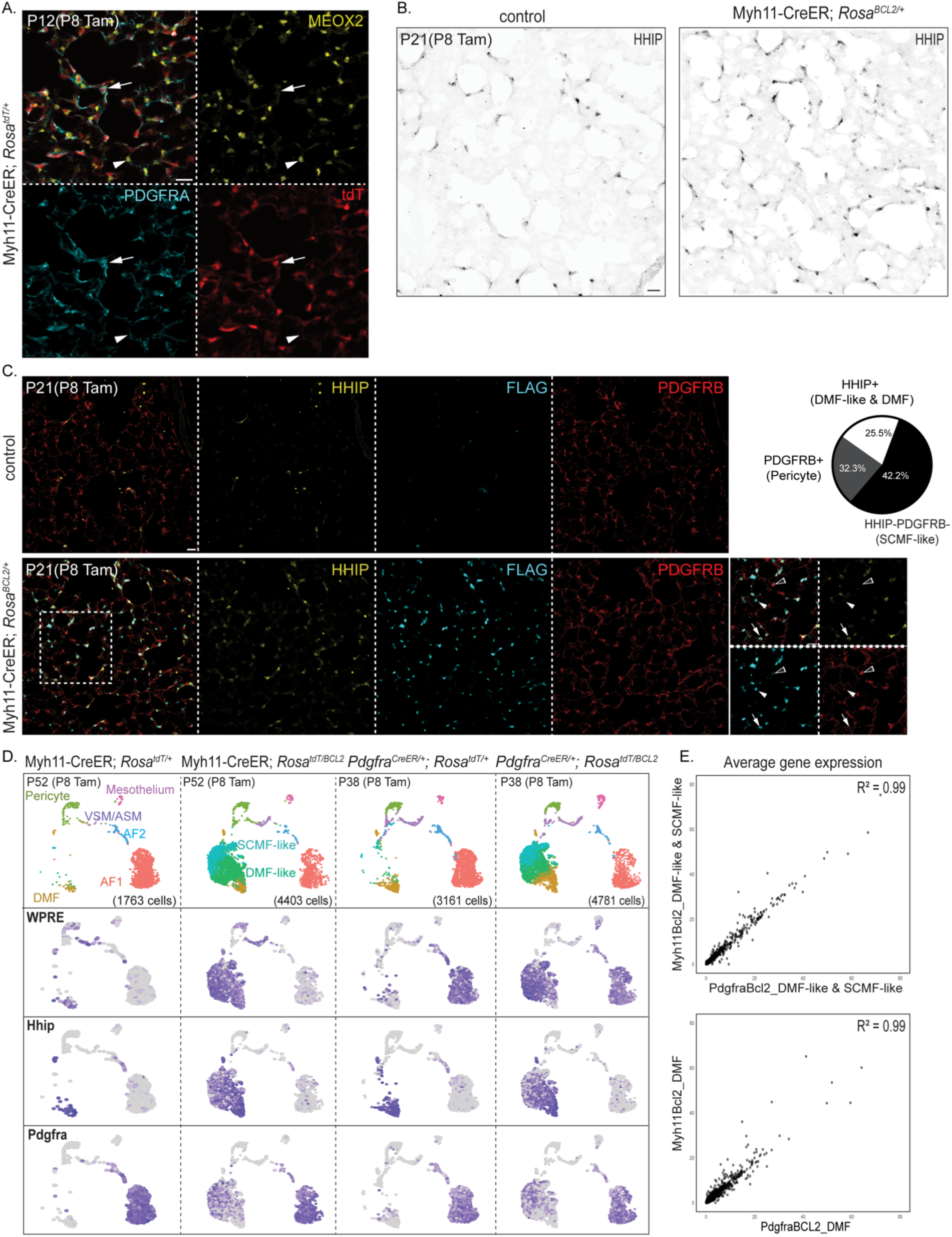
Myh11-CreER driver recapitulates the persistent AMF phenotype. A. Immunostaining of acutely labeled lungs, representative of at least 4 mice, showing that Myh11-CreER targets AMFs (PDGFRA^high^; arrow), but not AF1s (MEOX2+; arrowhead). B. Immunostaining of mature Myh11-CreER; *Rosa^BCL2/+^* and control lungs, representative of at least 4 mice, showing more HHIP+ cells in BCL2 lungs extending to distal alveolar ducts. C. Immunostaining and quantification of mature Myh11-CreER; *Rosa^BCL2/+^*and control lungs, showing the expansion of persistent populations FLAG+HHIP-PDGFRB-(SCMF-like; arrowhead) and FLAG+HHIP+ (DMF and DMF-like; arrow) in Myh11-CreER; *Rosa^Bcl2/+^* lungs. This driver also targets pericytes (PDGFRB+FLAG+; open arrowhead). D. Top: UMAPs of lung mesenchymal cells from adult Myh11-CreER; *Rosa^BCL2/+^* and *Pdgfra^CreE^*^R/+^*; Rosa^BCL2/+^* and their respective control littermates showing comparable persistent cells (SCMF-like and DMF-like) with both drivers. Bottom: Feature plots show that the only cells targeted by both drivers (WPRE+) are DMF-like and SCMF-like cells, but not AF1/2 cells, pericytes, or A/VSMCs. E. Scatterplot comparison of average gene expression of AMFs (including SCMF-like and DMF-like cells; top) and DMFs (bottom) from Myh11-CreER; *Rosa^BCL2/+^* and *Pdgfra^CreE^*^R/+^*; Rosa^BCL2/+^* lungs, showing a high correlation (R^2^=0.99) between drivers. All scale bars: 20 um.

To fully characterize BCL2-rescued cells in the Myh11-CreER model, we performed scRNA-seq on P52 Myh11-CreER; *Rosa^tdT/+^* control and Myh11-CreER; *Rosa^BCL2/+^* lungs. Only male mice were included since Myh11-CreER was Y-linked^49^. The WPRE lineage marker was expectedly restricted to VSMCs, ASMCs, DMFs, and pericytes, and not AF1s or AF2s, in the control but expanded to an ectopic population specific to the BCL2 lung (Figure 4D). We considered this population persistent AMFs, consisting of DMF-like and SCMF-like cells, as they aligned with the corresponding population in the *Pdgfra^CreER^* model and exhibited the same opposing gradients of *Hhip* and *Pdgfra* (Figure 4D). Additionally, persistent AMFs from both Myh11-CreER and *Pdgfra^CreER^*models exhibited a 99% correlation in gene expression, comparable to that observed for the BCL2-insensitive DMFs of each model (Figure 4E). Since the only population targeted by both Myh11-CreER and *Pdgfra^CreER^* was AMFs that underwent apoptotic clearance, the most plausible explanation for their shared phenotype was that BCL2-rescued AMFs were reprogrammed toward the DMF fate.

The third CreER driver, *Pdgfrb^CreER^*, was traditionally considered specific to VSMCs and pericytes^14,50^. However, we previously showed that it also targeted AF2s and, curiously, had ∼85% specificity despite achieving >95% efficiency in targeting pericytes in the alveolar region^14^. This non-specific labeling was ultimately identified as marking DMFs and AMFs, despite low PDGFRB expression in them (Figure 5A). In P55 lungs targeted at P8, the WPRE lineage marker labeled DMFs along with the expected VSMCs and pericytes in the control. In BCL2 lungs, however, WPRE expanded to the same ectopic population composed of DMF-like and SCMF-like cells, as supported by HHIP immunostaining (Figure 5B and 5C). The same high concordance in gene expression was observed for persistent AMFs from *Pdgfra^CreER^*and *Pdgfrb^CreER^* models (Figure 5D). Thus, our scRNA-seq characterization of three BCL2 models using *Pdgfra^CreER^*, Myh11-CreER, and *Pdgfrb^CreER^* clarifies driver specificity and collectively establishes AMFs as the origin of persistent cells.

**Figure 5.**
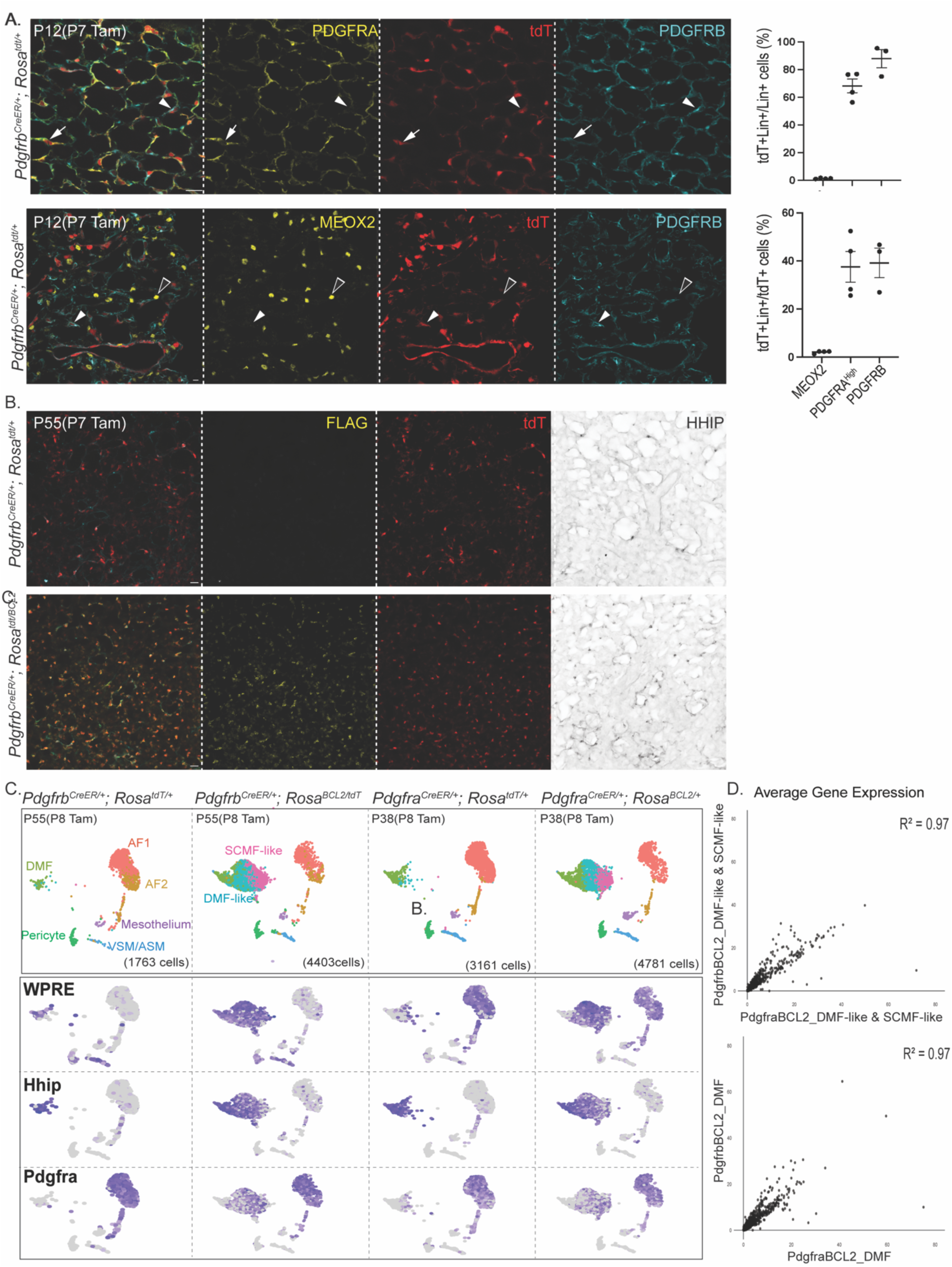
*Pdgfrb^CreER^* driver recapitulates the persistent AMF phenotype. A. Immunostaining and quantification of acutely labeled lungs showing that *Pdgfrb^CreER^* targets pericytes (PDGFRB+; arrowhead) and AMFs (PDGFRA^High^; arrow), but not AF1s (MEOX2+; open arrowhead). B. Immunostaining of adult *Pdgfrb^CreER/+^; Rosa^BCL2/+^* and control lungs shows more HHIP+ cells in BCL2 lungs in distal alveolar ducts. C. Top: UMAPs of lung mesenchymal cells from adult *Pdgfrb^CreE^*^R/+^*; Rosa^BCL2/+^* and *Pdgfra^CreE^*^R/+^*; Rosa^BCL2/+^*and their control littermates showing comparable persistent cells (SCMF-like and DMF-like) with both drivers. Bottom: Feature plots showing opposing gradients of Pdgfra and Hhip in targeted (WPRE+) SCMF-like and DMF-like cells, respectively, in both drivers. D. Scatterplot comparison of average gene expression of AMFs (including SCMF-like and DMF-like cells; top) and DMFs (bottom) from *Pdgfrb^CreE^*^R/+^*; Rosa^BCL2/+^* and *Pdgfra^CreE^*^R/+^*; Rosa^BCL2/+^*, showing a high correlation (R^2^=0.97) between drivers. All scale bars: 20 um.

### Both DMF and DMF-like cells upregulate contractile genes upon house dust mite (HDM) exposure

To investigate the functional implication of persistent AMFs, we examined whether transcriptomic similarity between DMFs and DMF-like cells translated into pathophysiological convergence, particularly in response to lung injury. Little was known about the role of DMFs beyond their proposed structural function in shaping alveolar ducts^14,26,43^. Since mesenchymal cells of the epithelial axis could share functions and ASMCs were known to respond to allergic inflammation in asthma^51–53^, we hypothesized that DMFs might respond as well. Indeed, in a 3-week HDM model, DMFs that had previously downregulated contractile genes upon the completion of neonatal alveologenesis reactivated them (Figure 6A). This response was exclusive to DMFs, as increased contractile gene expression was confined to HHIP+ cells along the proximal alveolar ducts (Figure 6A). In BCL2 lungs from the Myh11-CreER model, contractile gene-expressing cells extended to distal alveolar ducts and were HHIP+tdT+, thus corresponding to DMF-like cells (Figure 6B). In contrast, neither SCMF-like cells nor pericytes (HHIP−tdT+) had this response (Figure 6C). These findings uncover a previously unrecognized response of DMFs to allergic inflammation and highlight a pathophysiological convergence between DMFs and persistent AMFs in this context.

**Figure 6.**
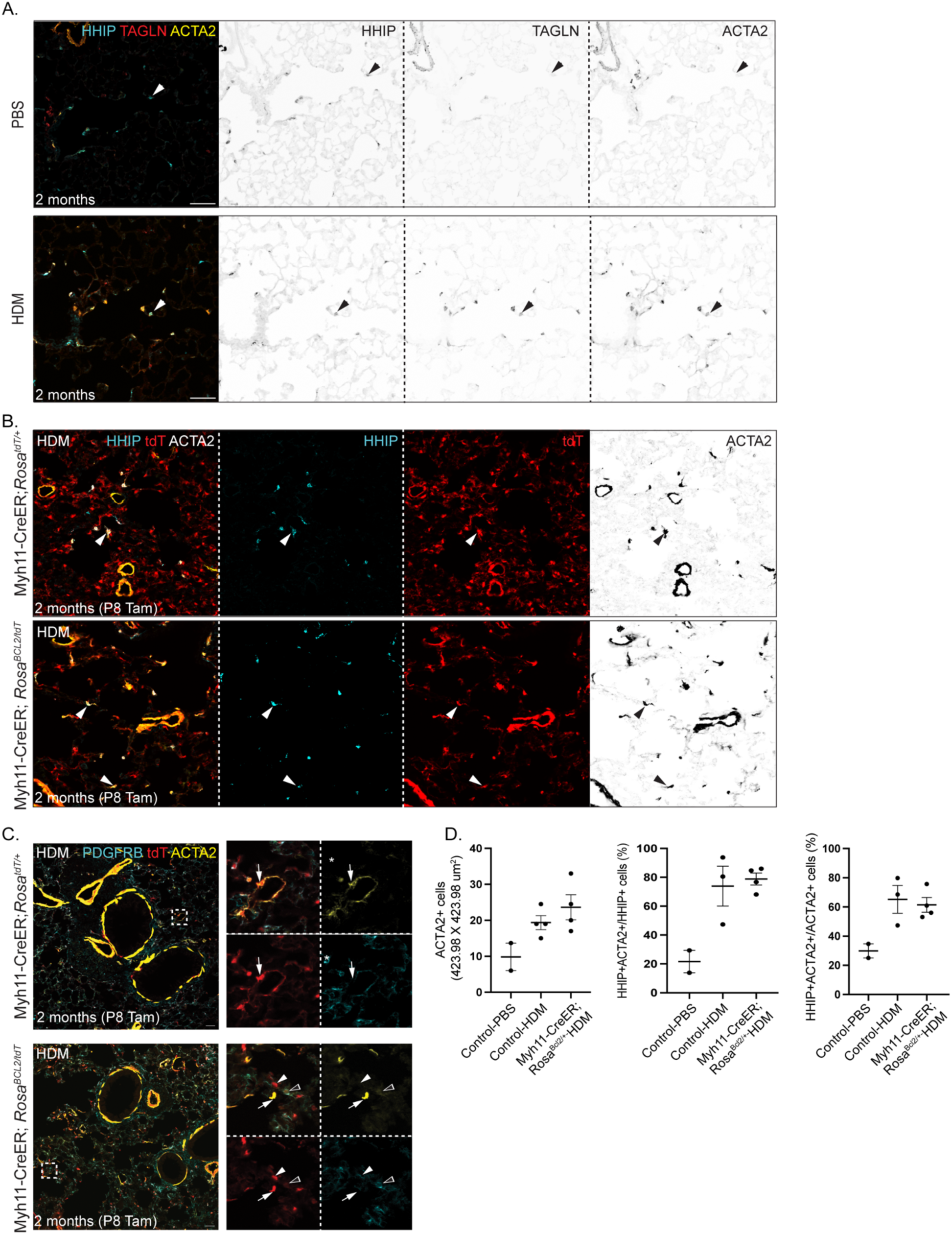
HDM exposure reactivates contractile genes in DMFs and DMF-like cells. A. Immunostained adult lungs exposed to 3 weeks of HDM showing upregulation of ACTA2 and TAGLN specifically in DMF (HHIP+; arrowhead), in comparison to PBS exposure. B. Immunostained adult control and Myh11-CreER; *Rosa^BCL2/tdT^* lungs exposed to 3 weeks of HDM showing that BCL2 lungs have an expansion of ACTA2+ cells to distal alveolar ducts, corresponding to DMF-like cells (HHIP+tdT+; comparing arrows in control and BCL2 lungs). C. Immunostained adult control and Myh11-CreER; *Rosa^BCL2/tdT^* lungs exposed to 3 weeks of HDM showing other lineage-labeled cells including SCMF-like cells (PDGFRB-tdT+; arrowhead) and pericytes (PDGFRB+tdT+; open arrowhead) do not express ACTA2. Arrow: ACTA2+tdT+ DMFs or DMF-like cells. D. Quantification showing comparable increase in ACTA2+ cells and the percentages of HHIP+ACTA2+ cells within HHIP+ or ACTA2+ alveolar cells in control and Myh11-CreER; *Rosa^BCL2/tdT^* lungs upon HDM exposure, in comparison to PBS exposure. Note that BCL2 lungs have more HHIP+ cells. All scale bars: 20 um.

## DISCUSSION

In this study, we have generated an inducible BCL2 overexpression allele that can be activated intersectionally with Flp and Cre, facilitating apoptosis research by shifting the balance of pro-survival and pro-apoptosis factors to inhibit apoptosis. Our BCL2 model phenocopies classical developmental apoptosis mutants of pro-apoptosis genes such as Bax, Bak, and Bok^27,31^. Using three independent Cre drivers that overlap only for lung AMFs, we provide causal evidence for AMF apoptosis and uncover the unexpected plasticity of rescued AMFs to mimic DMFs molecularly and functionally in response to HDM-induced allergic inflammation (Figure 7). Our BCL2 model and insights into lung mesenchymal cells of the epithelial axis advance the field of lung mesenchymal development and diseases.

**Figure 7.**
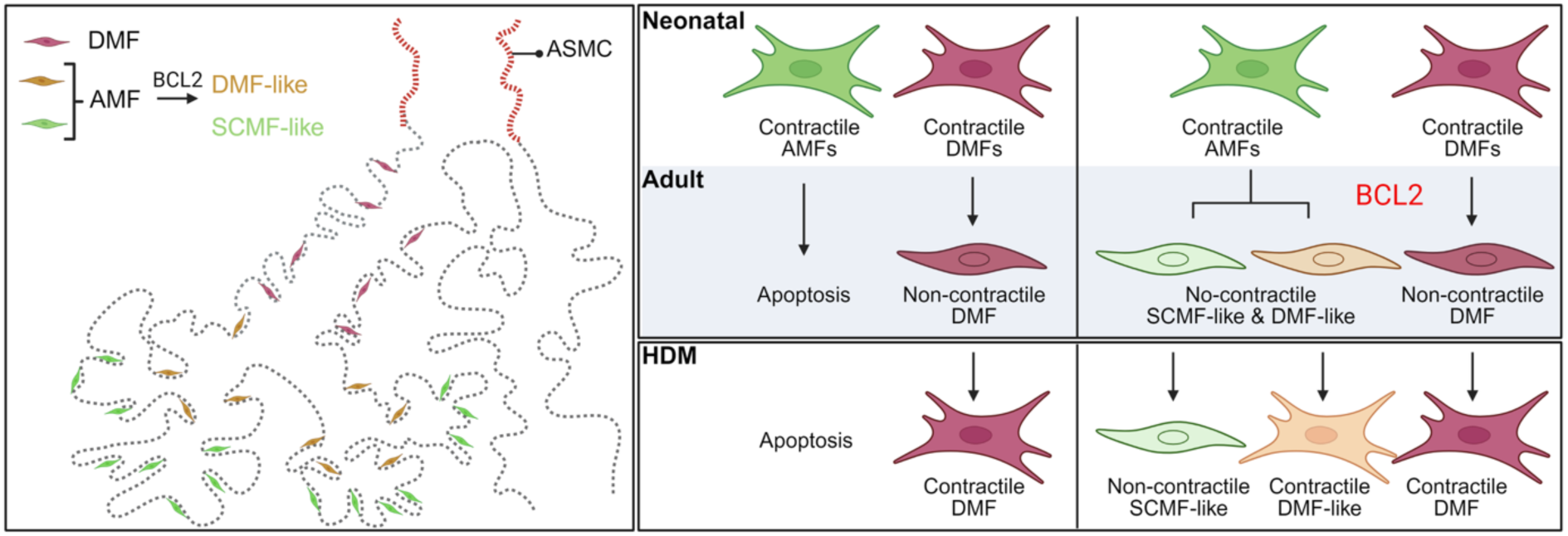
Diagram summary of lung mesenchymal cells of the epithelial axis and their responses to BCL2 overexpression and HDM exposure. Left: Schematics with color-coded cell types showing, from proximal to distal, ASMCs, DMFs and AMFs surrounding the lung epithelium. DMFs surround proximal alveolar ducts, while AMFs surround distal alveolar ducts and alveoli, and, upon BCL2 overexpression, become DMF-like (HHIP+) and SCMF-like (HHIP-) cells. Right: Neonatal AMFs and DMFs are contractile, and undergo apoptosis or become non-contractile, respectively. Upon HDM exposure, adult DMFs reactivate contractile genes, so do persistent DMF-like cells, but not persistent SCMF-like cells.

Studying apoptosis in vivo is akin to catching a falling knife, as only a small fraction of dying cells are present at any given time and often deteriorate too quickly to retain cell type markers. The human BCL2 protein in our inducible model functions across multiple mouse tissues, offering a versatile tool to prevent specific apoptotic events. Unlike other lung cells, AMFs are a transient population, undergoing apoptosis potentially to minimize cellular components at the gas exchange interface after shaping the alveoli. AMF apoptosis may be triggered intrinsically by contraction-induced exhaustion or extrinsically from deprivation of physicochemical survival signals from the extracellular matrix or neighboring cells. Regardless, BCL2 overexpression, but not endogenous BCL2, is sufficient to inhibit AMF apoptosis (Figure 1). Identifying pro-apoptotic proteins that BCL2 counteracts, such as BAX, BAK, and/or BOK, and unraveling their upstream regulators will provide further insights into AMF apoptosis mechanisms, which may be disrupted during impaired alveologenesis in bronchopulmonary dysplasia.

The molecular and functional convergence of rescued AMFs toward DMFs supports our three-axis classification of the heterogeneous lung mesenchymal cells. Just as SOX9 epithelial progenitors at branch tips leave behind progeny that differentiate into SOX2+ airway cells early in development and alveolar cells later, mesenchymal progenitors ahead of branch tips may leave behind progeny that differentiate into ASMCs early in development, followed by DMFs and AMFs ^54–57^. The choice among ASMCs, DMFs, and AMFs may be determined by the associated epithelium and/or the duration of differentiation. Specifically, while ASMCs align with airway cells ending at bronchoalveolar duct junctions, DMFs may align with bronchovascular bundles ending at arteriole-capillary junctions. AMFs, marked by high PDGFRA and targeted by *Pdgfra^CreER^*, surround distal alveolar ducts, primary and secondary septa. Those around the distal alveolar ducts may be maturing into DMFs but are normally removed via apoptosis; however, with BCL2 rescue, they become DMF-like cells (Figure 7). In this context, cell death becomes the ultimate way to restrict plasticity and end development. Future epigenomics analysis, including dissection of enhancers and associated regulators of marker genes such as *Actc1*, *Hhip*, and *Pdgfra*, will provide a path to understanding cell fate and plasticity within the epithelial axis. Similarly, VSMCs and pericytes within the vascular axis, as well as AF1s and AF2s within the interstitial axis, may share developmental origins and cell fate control mechanisms.

The re-expression of contractile genes in DMFs and even BCL2-rescued DMF-like cells in response to HDM-induced allergic inflammation (Figure 7) is unexpected yet, in retrospect, aligns with the three-axis model, considering the well-established hyperplasia and hypertrophy response of ASMCs^51,53^. If the developmental contractility of DMFs serves to counterbalance alveolar duct expansion, it remains unclear whether mechanical forces from HDM-induced tissue remodeling or inflammatory signals from immune or structural cells initiate the same contractile program. It is conceivable that children of immature lungs with still contractile DMFs might be more susceptible to asthmatic triggers, highlighting disease relevance of lung maturation. Again, dissection and comparison of developmental versus pathological enhancers of contractile genes will provide insight into these mechanisms. Finally, it is intriguing to speculate that each mesenchymal cell axis contributes uniquely to different lung diseases: the epithelial axis to asthma, the vascular axis to pulmonary hypertension, and the interstitial axis to pulmonary fibrosis^12,18,19,25,58^.

## RESOURCE AVAILABILITY

### Lead contact

Further information and requests for resources should be directed to and will be fulfilled by the lead contact, Jichao Chen (^28^).

### Materials availability

Mouse lines generated in this study are available from the lead contact upon request.

### Data availability statement

scRNA-seq data have been deposited in GEO under the accession number GSE297319.

## EXPERIMENTAL MODELS

### Generation of Rosa^BCL2+F^ knock-in mouse model

The targeting vector *Rosa^BCL2+F^*was generated by replacing the tdT reporter in the Ai65 targeting vector (Addgene #61577) with FLAG-tagged human BCL2 (Addgene #8768). Specifically, the FLAG-hBCL2 was PCR amplified with 5’-ACTACT<u>ACGCGT</u>GCCACC**ATG***GATTACAAGGATGACGACGATAAG*ATGGCGCACGC TGGGAGAAC (MluI site underlined; start codon in bold; FLAG tag italicized) and 5’-ATCATC<u>ACGCGT</u>TCACTTGTGGCCCAGATAGG (MluI site underlined) and cloned into the MluI site in Ai65 and verified by Sanger sequencing. The *Rosa^BCL2+F^*knock-in allele was generated using CRISPR targeting via standard pronuclear injection at the Genetically Engineered Mouse Facility at MD Anderson Cancer Center^59^. Specifically, 400 nM gRNA (Synthego), 200 nM Cas9 protein (E120020-250ug, Sigma), and 500 nM circular donor plasmid were mixed in the injection buffer (10 mM Tris pH 7.5, 0.1 mM EDTA). The gRNA targeted 5′ - ACTCCAGTCTTTCTAGAAGATGG with last 3 nucleotides being the protospacer adjacent motif (PAM; not included in gRNA). The FRT-flanked stop cassette was removed by crossing it with *Rosa^Flp^* (Jackson Laboratory 009086)^60^. The resulting Rosa^BCL2^ allele and the original Rosa^BCL2+F^ allele were genotyped for left (Rosa26L) and right (Rosa26R) homology arms using primers 5’-GGGCGTACTTGGCATATGAT and 5’-CGCCTAAAGAAGAGGCTGTG, and 5-TGCAGCAGCACGTGTTGACAATTA and 5-CCATTCTCAGTGGCTCAACA, and distinguishable with primers 5’-GCAACGTGCTGGTTATTGTGCTGT and 5’-GTGCGCCATCTTATCGTCGT.

### Mice (Mus musculus)

The following mouse strains were obtained from the Jackson Laboratory: *Pdgfra^CreER^* (032770)^61^, *Pdgfrb^CreER^* (030201)^62^, Myh11-CreER (019079)^49^, *Rosa^tdT^* (007914)^63^, Sox2-Cre (008454)^40^, and *Shh^Cre^* (005622)^38^.

Intraperitoneal injection of 500 ug of tamoxifen (T5648; Sigma) dissolved in corn oil (C8267, Sigma) at a concentration of 10 mg/ml was performed at postnatal day (P) 8 to induce recombination. The time of evaluation and the number of animals per group in each experiment are specified in the figure legends. Unless specified, females and males were used, and littermates of the same sex were randomly assigned to experimental groups. All animal experiments were reviewed and approved by the Institutional Animal Care and Use Committee at MD Anderson Cancer Center and the University of Texas Health Science Center.

## METHOD DETAILS

### Antibodies

The following antibodies were used for immunostaining: rabbit anti-MEOX2 (1:200, NBP2-30647, Novus Biological), goat anti-HHIP (1:500, AF1568, R&D), rabbit anti-FLAG (1:500, ab205606, Abcam), rat anti FLAG-DYKDDDDK (1:500, NBP1-06712, Novus Biological), rat anti-CADHERIN 4 (1:20, MRCD5, Developmental Studies Hybridoma), goat anti-PDGFRA (1:1000, AF1062, R&D); Alexa Fluor 633 hydrazide (1:1000 of 2.5 mg/ml stock dissolved in PBS, A30634, Thermo Fisher), rat anti-PDGFRA, (1:1000, 14-1401-82, eBioscience), goat anti-PDGFRB (1:1000, AF1042, R&D systems), rat anti-PDGFRB (1:1000 R&D systems AF2419), rabbit anti-TAGLN (1:1000, ab14106, Abcam), Cy3-conjugated mouse anti-Smooth muscle actin (1:1000, C6198, Sigma Aldrich), Alexa 647-conjugated mouse anti-Smooth muscle actin (1:1000, sc-32251 AF647, Santa Cruz Biotechnology), rabbit anti-cleaved Caspase-3 (1:500, 9661, Cell signaling), rabbit anti-TOMM20 (1:500, GTX32928, Genetex), goat anti-NOTCH3 (1:500, AF1318, R&D) rabbit anti-AQP5 (1:1000, ab78486, Abcam), rabbit anti-NK homeobox 2-1 (NKX2-1,1:1000, sc-13040, Santa Cruz). Alexa 488-, Cy3- or Alexa 647-conjugated donkey anti-rabbit (711-545-152, 711-165-152, 711-605-152), anti-rat (712-545-153, 712-165-153, 712-605-153), anti-goat (705-545-147, 705-165-147, 705-605-147) secondary antibodies (Jackson ImmunoResearch) were used.

### Harvesting lungs for immunostaining

Mice were anesthetized using Avertin (^4^, Sigma) by intraperitoneal injection. The right ventricle of the heart was then injected with phosphate-buffered saline (PBS, pH 7.4) to perfuse the lung. Then, the mice were cannulated, and the lungs were inflated with a solution of 0.5% Paraformaldehyde (PFA, P6148, Sigma) in PBS under a constant pressure of 25 cm H_2_O. Inflated lungs were fixed in 0.5% PFA in PBS for 3 hours at room temperature then washed with PBS overnight at 4 °C on a rocker.

### Cryosection immunostaining

For cryopreservation, lung lobes were dissected and transferred to a 20% sucrose (S5-500, Fisher Scientific) in PBS solution containing 10% optimal cutting temperature compound (OCT, 4583, Tissue-Tek) to cryoprotect the samples. Samples were incubated overnight and frozen the next day in OCT at -80°C. After sectioning (5-20 µm thickness), the sections were blocked for 1 hour in PBS with 0.3% Triton X-100 and 5% normal donkey serum (017–000-121, Jackson ImmunoResearch), and then incubated in primary antibodies overnight at 4 °C. The next day, the sections were washed with PBS for 1 hour and then incubated with the secondary antibodies for another hour at room temperature. After incubation, the sections were washed with PBS as described before. Finally, the sections were mounted with Aquamount (18606, Polysciences).

### Wholemount immunostaining

This is based on a protocol published previously with minor modifications^64^. Around 3 mm wide strips from the outer edge of the cranial or left lobe were cut and then blocked with PBS with 0.3% Triton X-100 (PBST) and 5% normal donkey serum. Then, the strips were incubated with primary antibodies diluted in PBST overnight at 4 °C. The next day, the strips were washed at room temperature with PBS+1% Triton X-100+1% Tween-20 (PBSTT) on a rocker for one hour. This step was repeated 3 times. Secondary antibodies and DAPI diluted in PBST were added and incubated overnight at 4 °C. After incubation, the strips were washed as described above and then fixed with 2% PFA in PBS for at least 2 hours. Finally, the strips were washed with PBS and mounted on slides using Aquamount (18606, Polysciences).

### Confocal microscopy and image analysis

An Olympus FV1000 confocal microscope was used for image acquisition. For wholemount immunostaining, Z-stack images of 20–30 μm thick at 1 µm-2 µm step size, and for cryosections, Z-stack images of 10-20 μm thick at 0.8 µm-1 µm step size were taken. Imaris 9.9 (Bitplane) was used for image visualization and analysis.

### Tissue dissociation and fluorescence-activated cell sorting

Adult lungs were dissected, minced into small pieces and digested at 37°C for 45 min in 1.35 ml Liebovitz media (Gibco, 21083–027) with the following enzymes: 2 mg/ml collagenase type I (Worthington, CLS-1, LS004197), 0.5 mg/ml DNase I (Worthington, D, LS002007), and 2 mg/ml elastase (Worthington, ESL, LS002294). 300 μl of fetal bovine serum (FBS, Invitrogen, 10082–139) was added to stop the enzymatic reaction. The tissue was mechanically dissociated by repeatedly pipetting the tissue up and down and filtered through a 70 μm cell strainer (Falcon, 352350). In a cold room, the samples were spun down at 1680 rcf for 1 min. The cells were resuspended in red blood cell lysis buffer (15 mM NH_4_Cl, 12 mM NaHCO_3_, 0.1 mM EDTA, pH 8.0) and incubated for 3 min, washed and resuspended with Liebovitz + 10% FBS. The cells were then filtered through a 35 µm cell strainer into a 5 ml glass tube and incubated for 30 minutes in ECAD-488 (53-3249-80, eBioscience), CD45-PE/Cy7 (103114, BioLegend and ICAM2-A647 (A15452, Invitrogen) at a concentration of 1:250. Cells were washed and resuspended with Liebovitz + 10% FBS and filtered again through a 35 µm strainer into a 5 ml glass tube. Sytox Blue was added as a viability dye (1:1000, Invitrogen, S34857). Cells were sorted using Aria II Cell sorter and FlowJo 10 was used to analyze the data. Dead cells and doublets were excluded and four cell lineages, immune cells (CD45^+)^, epithelial cells (ECAD^+^), endothelial cells (ICAM2^+^), and mesenchymal cells (CD45^-^ Ecad^-^ICAM2^-^) were collected for scRNA-seq.

### Single-cell RNA-sequencing

FACS-purified lung cells for each sample were combined as follows, 50% mesenchymal, 16.6% epithelial, 16.6% endothelial, and 16.6% immune cells. The cells were then processed through the Chromium Single Cell Gene Expression Solution Platform (10x Genomics) and partitioned using the Chromium Single Cell 3’ Library and Gel Bead Kit following the manufacturer’s user guide (v2, rev D). Samples were sequenced with an Illumina NextSeq500 using a 26X124 sequencing run format with an 8 bp index (Read1). scRNA-seq data output was pre-processed using CellRanger and the mouse reference genome GRCm38 (mm10) was used for read alignment. Further analysis was carried out using the R package Seurat 5.0. Detailed R script of the analysis in Supplemental File 1. Selected cells were filtered based on the following parameters: at least 200 detected genes but no more than 5000 genes and no more than 15% mitochondrial genes. Epithelial, immune, endothelial, lineages were identified based on the expression of *Cdh1*, *Ptprc*, *Icam2* respectively. *Col3a1* and *Col1a1* were used to identify mesenchymal cells. Doublets were filtered out and mesenchymal cells were subset and re-clustered. Findmarkers were used to identify differential genes between clusters and the R package Enhanced Volcano 1.0.1. was used for plotting with a fold change cutoff of 1 and a p-value cut off at 10e-9 unless otherwise stated in the figure legend. R Package Enrichr 3.2^65^ was used for gene ontology analysis, CellChat (1.6.1)^66^ for ligand-receptor analysis, and Monocle2^44^ for trajectory analysis. Publicly available datasets were obtained from the Gene Expression Omnibus (GEO) database GSE180822 ^14^ including ages P7, P13, and P20 were analyzed as described previously and integrated using IntegrateData.

### Lung function analysis

Adult mice were weighed and anesthetized with an intraperitoneal dose of Avertin (240 mg/kg). Then each mouse was subjected to tracheotomy and cannulated with a blunted 18-gauge needle, and ventilated mechanically using the FlexiVent System (SCIREQ, Montreal, Canada). The mechanical scan program was used and included: snapshot to measure tracheal dynamic resistance, dynamic compliance, and dynamic elastance; Quick Prime that measures Newtonian resistance, airway resistance (Rn), tissue damping (G), and the tissue elastance (H); Pressure-volume (PV) loop: the quasi-static compliance (Cst) and quasi-static elastance (Est), pressure-volume curves, estimate of inspiratory capacity (A), curvature (K), and area of the PV loop (Hysteresis).

House dust mite (HDM)-induced allergic asthma

Wild type, Myh11-CreER; *Rosa^tdT/+^* and Myh11-CreER; *Rosa^BCL2/tdT^* mice at least 8 weeks of age were anesthetized with isofluorane and suspended by the maxillary incisors on a board with a 60° incline. Mice were treated with 15 ug HDM (NC9756554, Greer Laboratories) in 50 ul PBS through oropharyngeal aspiration every other day for three weeks. Mice were euthanized for analysis after 24 hours of the last dose. Mice in the control group were instilled with 50 μL PBS.

## QUANTIFICATION AND STATISTICAL ANALYSIS

### Mean Linear Intercept

The mean linear intercept measurements were performed on 5 um-thick fixed-frozen lung sections stained with hematoxylin and eosin. For each lung, four images were acquired per sample on an upright Olympus BX60 microscope with a 20X objective. Horizontal and vertical gridlines were drawn evenly spaced and the distance from one alveolar wall to the next along the gridlines was measured with Image J. Airways and main vessels were excluded.

### Elastic fiber analysis (Alexa-647 hydrazide area)

Surface objects were created using Imaris 9.9 and statistics for volume, and area measurements were calculated. Identical parameters were used between images and experiments. Manual editing for aberrant surfaces was performed before statistical analysis.

### Statistical analysis

All statistical analyses were performed in GraphPad Prism 9.0 using a two-tailed Mann-Whitney test unless otherwise indicated in figure legends. Results were considered significant if the p-value was < 0.05. The number of biological replicates were listed in figure legends.

## ACKNOWLEDGEMENTS

We thank the University of Texas MD Anderson Genetically Engineered Mouse Facility for generating the *Rosa^BCL2+F^*mice, with the support of the Cancer Center Support Grant (CA #16672). We thank Drs. Guolun Wang, Eric Brandt, Neeru Hershey, and Timothy Le Cras for pilot HDM experiments. ChatGPT was used to edit the manuscript. This work was supported by the University of Texas MD Anderson Cancer Center Retention Fund, National Institutes of Health R01HL130129, R01HL153511, R35HL171346 (JC) and Larry Deaven PhD Fellowship in Biomedical Sciences 2022-2023 (MJGG).

## AUTHOR CONTRIBUTIONS

MJGG and JC designed research; MJGG, HL performed research and analyzed data; JC generated the *Rosa^BCL2+F^*mice; MJGG, TMW, SEE, and JC wrote and edited the paper; all authors read and approved the paper.

---

**Supplemental Fig. 1.**
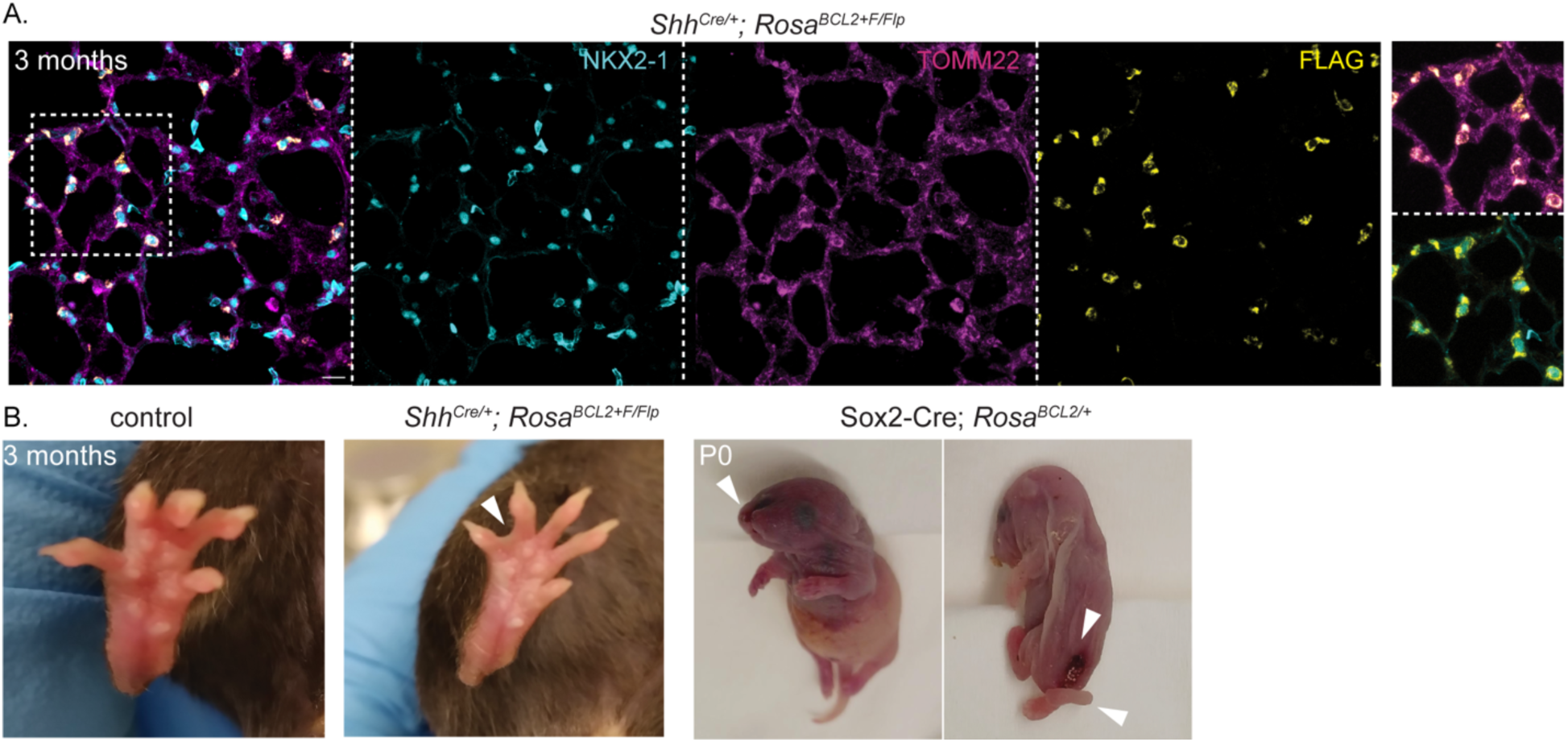
Mitochondrial localization and function of overexpressed BCL2. A. Immunostaining of adult lungs shows that exogenous FLAG-tagged BCL2 activated by *Shh^Cre^* is specific to epithelial cells (NKX2-1+) and colocalizes with a mitochondrial marker TOMM22. Scale bar: 20 um. B. Webbing between digits four and five (arrowhead) from exogenous BCL2 activated by *Shh^Cre^*. C. Developmental defects including cleft palate (left), sacral spina bifida, and curly tail (right) from whole-embryo exogenous BCL2 activated by Sox2-Cre.

**Supplemental Fig. 2.**
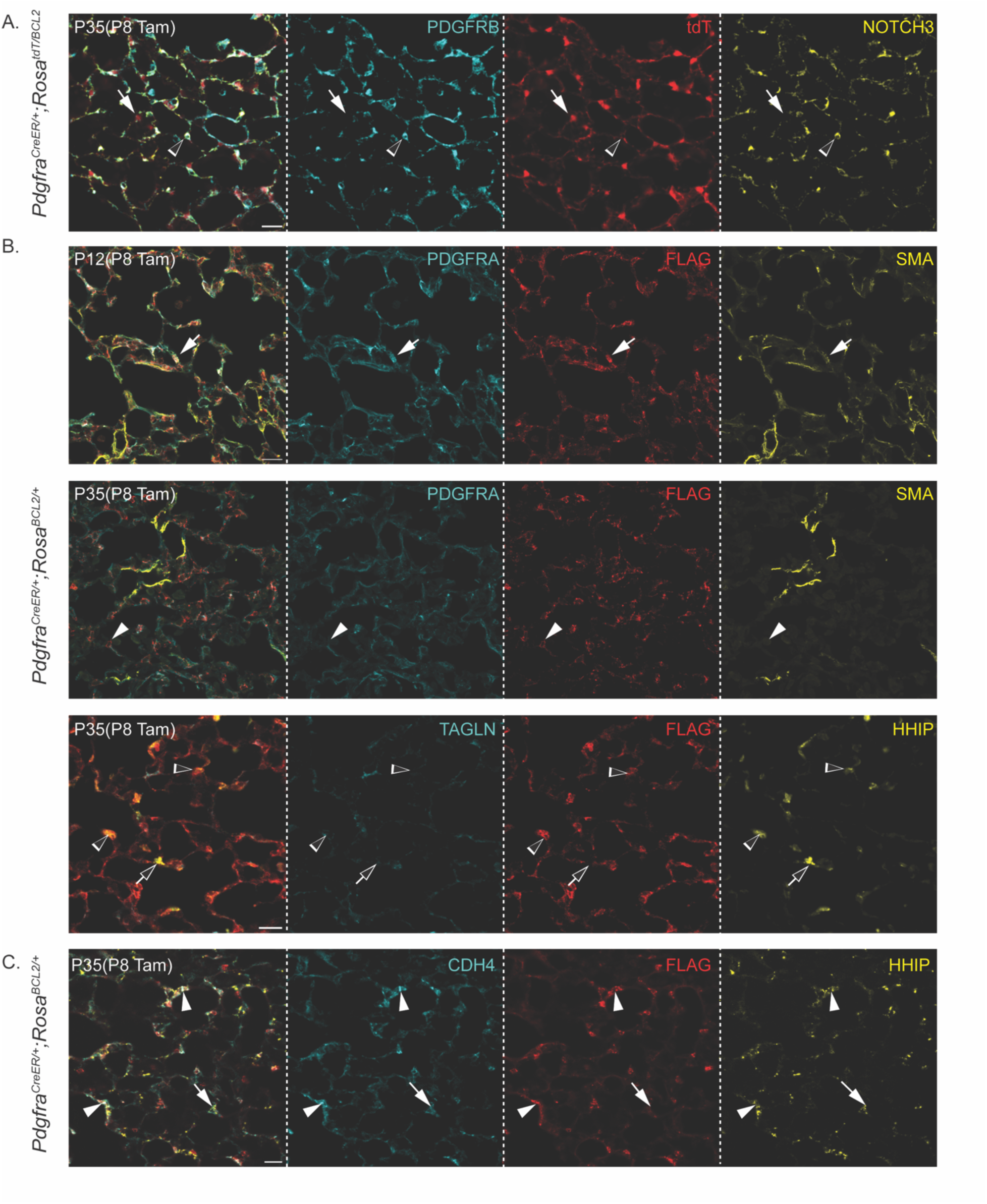
Persistent AMFs remain in the epithelial axis and downregulate contractile proteins. A. Immunostaining of mature lungs, representative of at least 4 mice (same below), shows that persistent cells (arrow) do not express PDGFRB or NOTCH3, markers of the vascular axis (open arrowhead). B. Immunostaining of neonatal and mature BCL2 lungs shows that neonatal AMFs (FLAG+PDGFRA^high^; arrow in top row) are ACTA2+, while persistent AMFs (FLAG+; arrowhead in middle row) downregulate PDGFRA and ACTA2. Bottom row: DMF-like cells (FLAG+HHIP+; open arrowhead) and DMF (FLAG-HHIP+; open arrow) cells also downregulate TAGLN. C. Immunostaining of mature BCL2 lungs shows that DMF-like cells (FLAG+; arrowhead) co-express HHIP and CDH4, markers of DMFs (FLAG-; arrow). All scale bars: 20 um.

**Supplemental Fig. 3.**
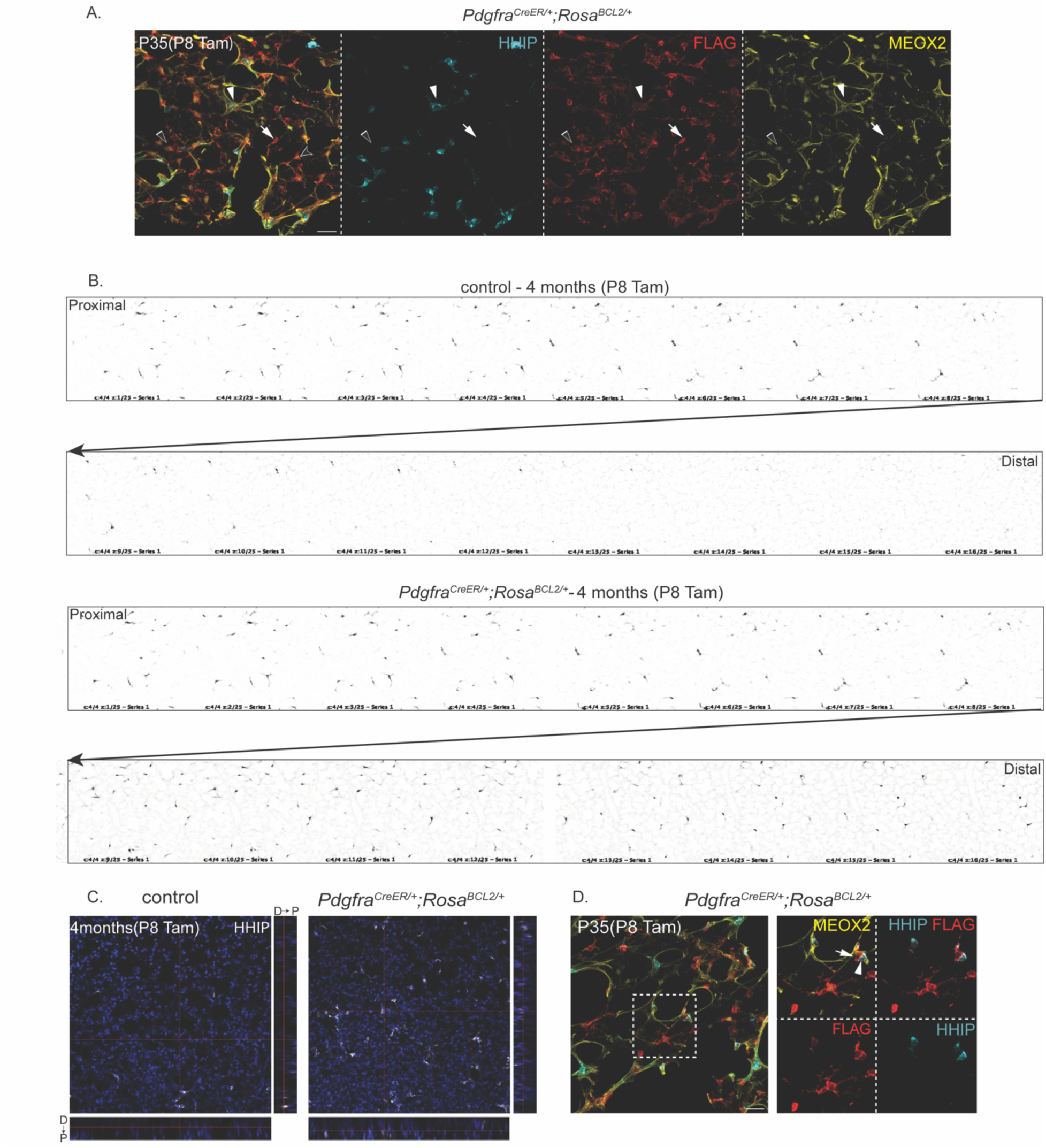
DMF-like cells surround distal alveolar ducts while SCMF-like cells surround alveoli. A. Immunostaining of mature BCL2 lung shows DMF-like cells (FLAG+HHIP+; arrowhead) surrounding distal alveolar ducts, while SCMF-like cells (FLAG+HHIP-; arrow) reside in the terminal alveoli. Like DMFs, DMF-like cells are associated with thick elastin fibers (autofluorescence in MEOX2 staining), whereas SCMF-like cells are associated with none or thin elastic fibers. Open arrowhead: FLAG+MEOX2+ AF1s targeted by *Pdgfra^CreER^*. B. Single-section montages of Z stacks of wholemount immunostained lung strips shows that DMFs (HHIP+) in the control are restricted to proximal alveolar ducts, whereas DMF-like cells (HHIP+) in the BCL2 lung expand distally. C. Orthogonal views of Z stacks of wholemount immunostained lung strips show more HHIP+ cells located distally in the BCL2 lung than the control. D: distal; P: proximal. D. Immunostaining of mature BCL2 lungs showing seemingly juxtaposition of persistent SCMF-like (arrow) and DMF-like (arrowhead) cells due to the 3D tissue complexity. All scale bars: 20 um.

**Supplemental Fig. 4.**
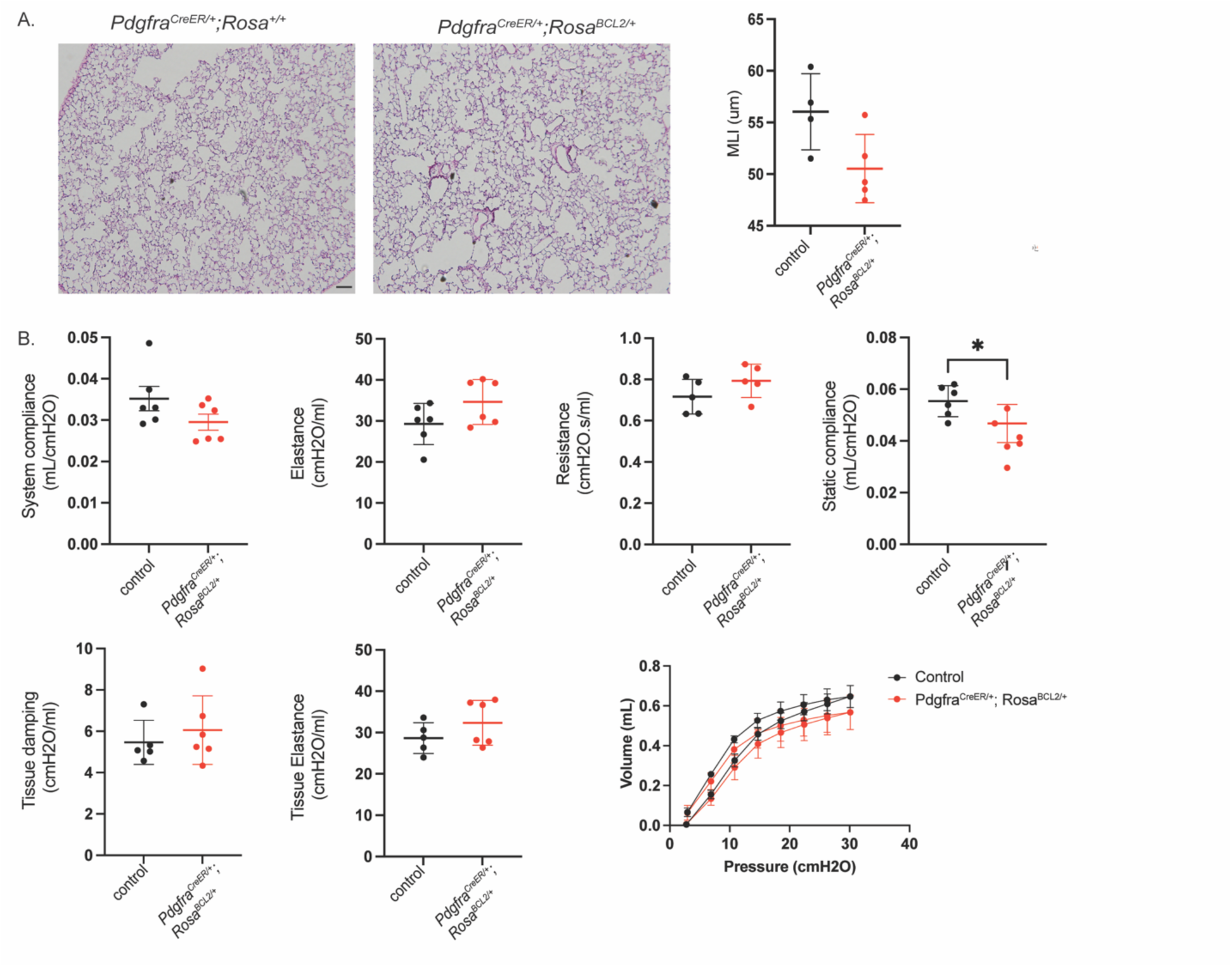
Persistent AMFs minimally affect lung morphology or mechanics. A. H&E staining of BCL2 and littermate control lungs and mean linear intercept (MLI) quantification show no significant difference (n=4; 3 images/mouse). Scale bar: 20 um. B. FlexiVent measurements of lung mechanics show minimal change in BCL2 lungs except for a small decrease in static compliance. See Table S1 for raw data.

**Supplemental 5.**
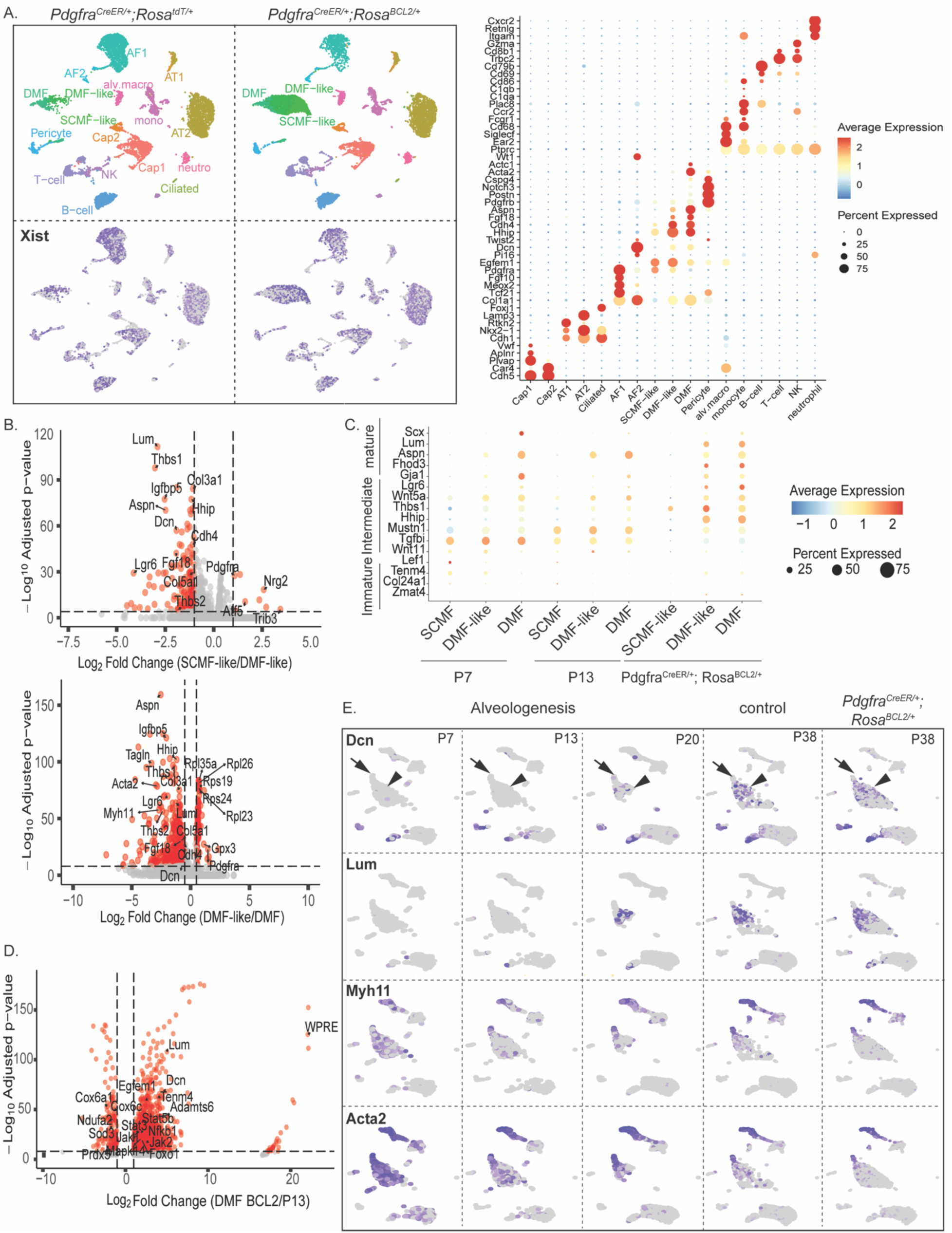
Persistent AMFs change from contractile to non-contractile fibroblasts. A. UMAP and dot plot of mature lungs color-coded by cell types of epithelial, endothelial, mesenchymal, and immune lineages (top) and feature plots of Xist showing comparable results from male and female lungs (bottom). B. Volcano plots comparison of SCMF-like and DMF-like (Top), and DMF-like and DMFs (Bottom) from mature BCL2 lungs, showing upregulation of genes associated with stages of myofibroblasts maturation. Genes such as *Pdgfra* and ribosomal genes (*Rpl28*, *Rpl23*, *Rps19*) found in SCMF-like and DMF-like represent immature and intermediate DMFs, while upregulation of DMF markers such as *Fgf18*, *Thbs2*, *Myh11*, and *Lum* corresponds to more mature DMFs. C. Dot plots show that previously reported immature (*Zmat4*, *Col24a1*,*Tenm4*, *Lef1*) intermediate (*Mustn1*, *Hhip*, *Thbs1*, *Wnt5a*), and mature (*Scx*, *Lum*, *Aspn*, *Fhod3*) myofibroblast markers of the developing lungs (P7 and P13) generally align with SCMF-like cells, DMF-like cells, and DMFs in BCL2 lungs. D. Volcano plot comparison of DMFs from P13 control and mature BCL2 lungs showing upregulation of mature DMF genes (*Lum* and *Dcn*) and genes associated with survival and differentiation (*Jak1*, *Jak2*, *Stat3*) in BCL2 lungs. Upregulation of the lineage marker WPRE suggests that some persistent cells are reprogrammed completely into DMFs. E. Feature plots showing that like DMFs (arrow), DMF-like cells (arrowhead) mature by upregulating matrix genes Dcn and Lum and downregulating contractile genes Myh11 and Acta2.

**Supplemental 6.**
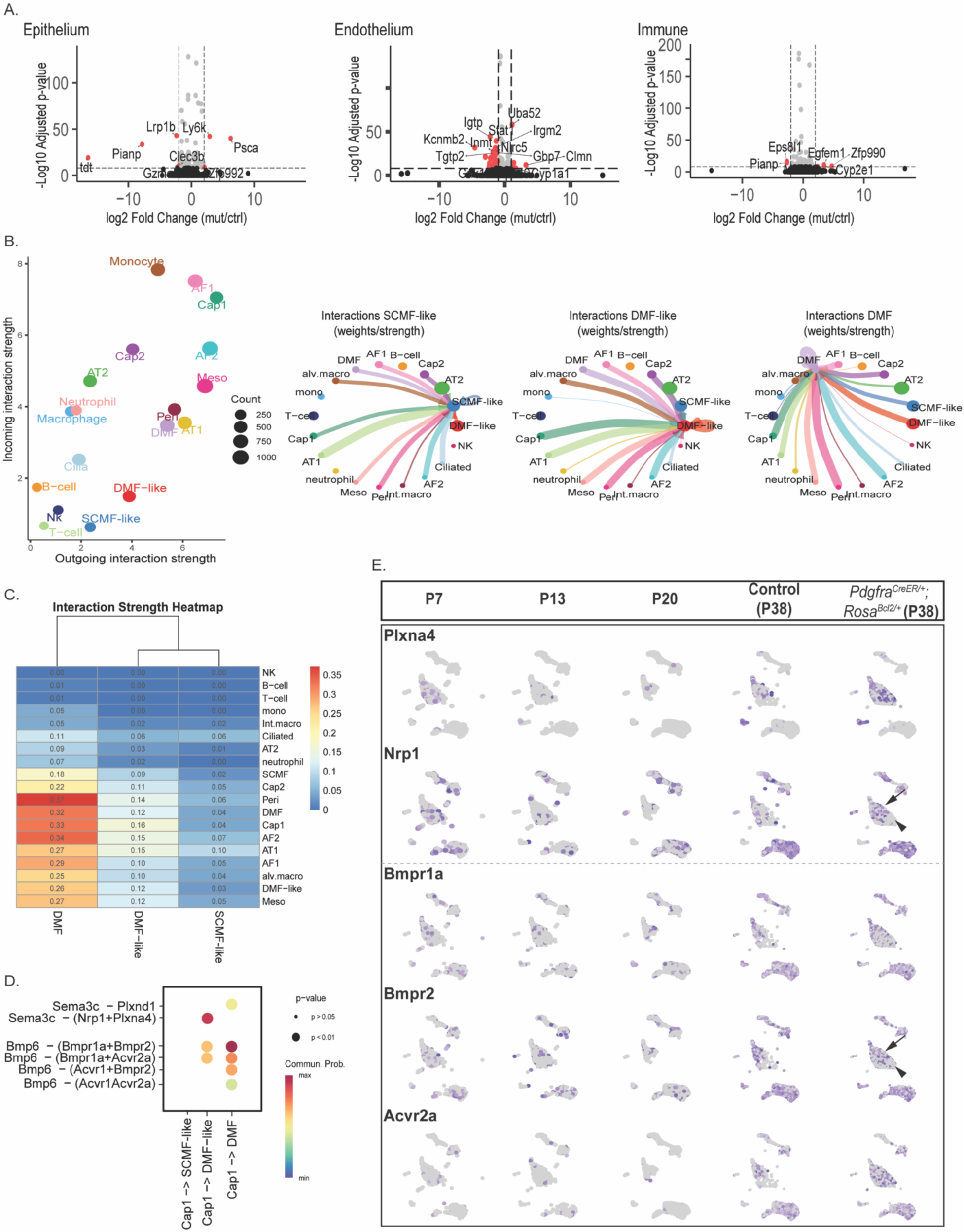
Limited interactions between persistent AMFs and other cell lineages. A. Volcano plots comparison of epithelial, endothelial and immune lineage cells between P38 *Pdgfra^CreER/+^; Rosa^BCL2/+^* and control lungs showing minimal changes in non-mesenchymal cell lineages. B. Left: CellChat ligand-receptor analysis of BCL2 lungs showing low outgoing and incoming interaction strength of SCMF-like and DMF-like cells. Right: Circle plots showing increasing interaction weights/strength for SCMF-like cells, DMF-like cells, and DMFs. C. Heatmap of interaction strength showing strongest interactions between DMFs and CAP1s, DMFs, pericytes, and AF2s. D. Differential incoming ligand-receptor interactions (Semaphorin and Bmp) of DMF-like cells, but not SCMF-like cells, with CAP1s, possibly mediating DMF-like reprogramming. E. Feature plots of neonatal and mature lungs show that expression of potential Semaphorin (Nrp1) and Bmp (Bmpr2) receptors mediating DMF-like reprogramming is higher in DMF-like cells (arrow) than SCMF-like cells (arrowhead). Plxna4, a receptor for Sema3c/3d, is specific for mesenchymal cells of the epithelial axis.

**Supplemental 7.**
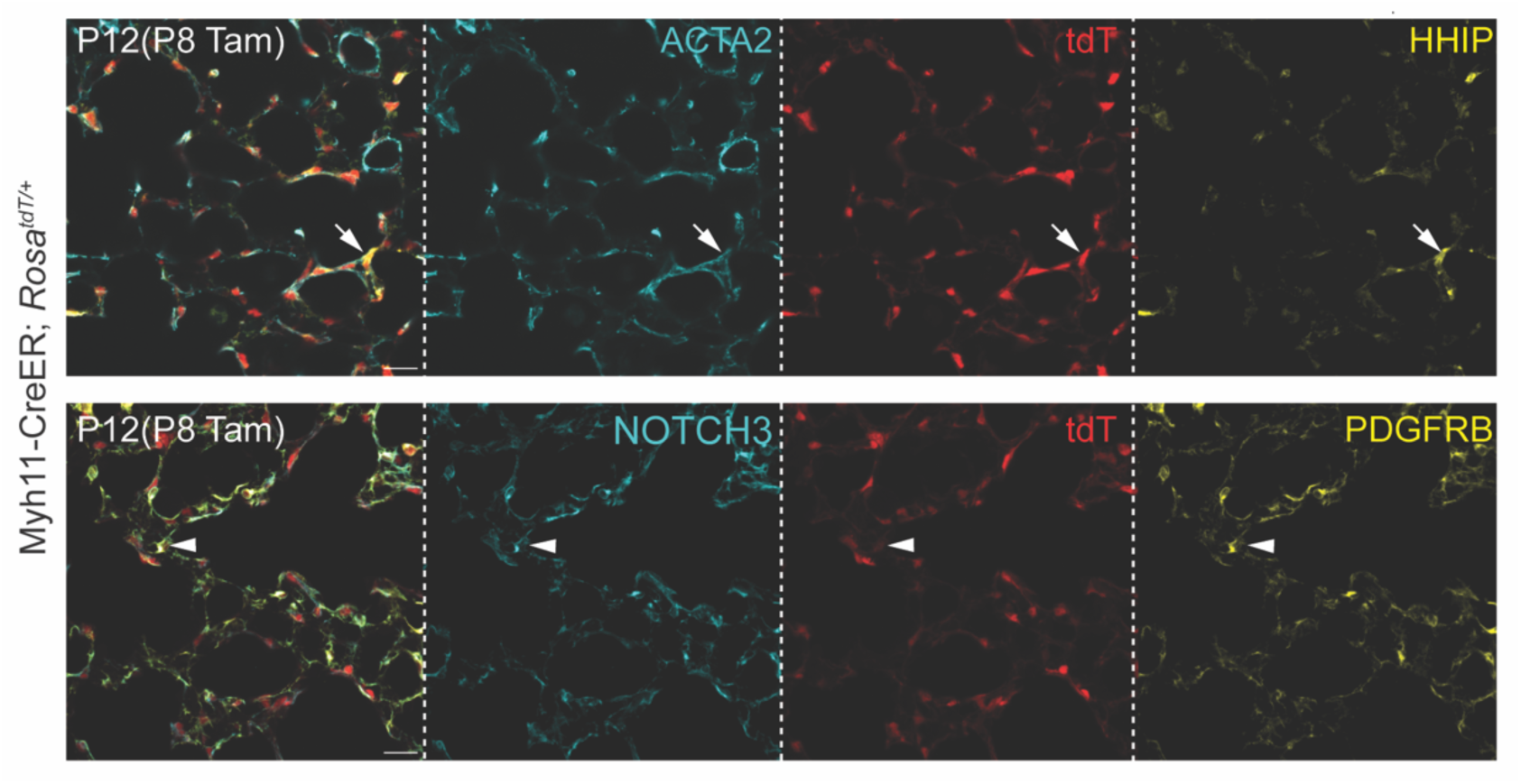
Additional characterization of the Myh11-CreER driver. Immunostaining of acutely labeled lungs, representative of at least 4 mice, showing that Myh11-creER targets DMFs (HHIP+; arrow) and pericytes (PDGFRB+NOTCH3+; arrowhead), although pericyte tdT is lower. Scale bar: 20 um.

**Supplemental Table S1: Raw data for image and flexiVent quantification.**

**Supplemental Table S2: Raw data for scRNA-seq plots.**

